# Circulating extracellular vesicles drive microglial senescence and neurodegeneration in Parkinson’s disease

**DOI:** 10.64898/2026.03.10.709299

**Authors:** Ankush Yadav, Elena Vacchi, Sandra Pinton, Edoardo Lazzarini, Matteo Pecoraro, Andrea Raimondi, Lucio Barile, Alain Kelin-Lang, Giorgia Melli

**Author notes:** Corresponding author and lead contact: Contact information, Giorgia Melli, Neurodegenerative Diseases Group, Institute for Translational Research, EOC-USI, Bellinzona, Switzerland.

## Abstract

**Background:** Extracellular vesicles (EV), secreted membrane particles involved in cell-to-cell communication, carry important information on immunity and its dysregulation. Recent studies have demonstrated the crucial role of peripheral and central inflammation in causing Parkinson’s disease (PD), as well as the involvement of EV in mediating neuron-glial interactions during neurodegeneration. However, the underlying mechanism of plasmatic EV in PD pathogenesis remains unknown.

**Methods:** EV were isolated from pool of plasma of PD patients and age- and sex-matched healthy controls (HC) using size-exclusion chromatography and characterized by nanoparticle tracking analysis, western blot, and transmission electron microscopy. SH-SY5Y neurons and HMC3 microglia cells were treated with EV, and their impact was evaluated using flow cytometry and immunofluorescence. Conditioned medium (CM) from EV-treated HMC3 cells was applied to SH-SY5Y neurons to determine indirect neurotoxic effects. Cytokine profiling and senescence-like features of EV-treated HMC3 cells were assessed. Unbiased proteomic analysis of PD-EV and HC-EV were further performed.

**Results:** PD-EV induced axonal degeneration and cell death in SH-SY5Y neurons and increased levels of TNF-α, IL-1β, IFN-γ, IL-8, and CCL11, accompanied by the expression of p16^INK4a^ in HMC3 cells, suggesting a proinflammatory, senescence-associated secretory phenotype (SASP). Enrichment pathway analysis revealed that these changes were mainly related to inflammatory and immune responses. Moreover, CM from PD-EV-HMC3 cells increased apoptotic cell death in SH-SY5Y neurons more than direct PD-EV. Notably, proteomic analysis of PD-EV showed higher expression of proteins involved in complement cascades, immune response, phagocytosis, and post translational protein translation, further supporting the potential of EV to induce inflammatory changes in PD.

**Conclusions:** This study demonstrates that plasmatic PD-EV contributes to neuronal degeneration by reducing neuronal integrity and indirectly by activating microglia through the secretion of pro-inflammatory, senescence-associated mediators. Circulating EV exerts a role in bridging peripheral inflammation with microglia, modulating neuroinflammatory events.

## Background

Parkinson’s disease (PD) is the second most common neurodegenerative disorder after Alzheimer’s disease, affecting approximately 7–10 million individuals worldwide^1,2^. The clinical presentation is dominated by motor symptoms such as rigidity, tremor, bradykinesia, and postural instability^3,4^. Neuropathologically, PD is defined by the progressive loss of dopaminergic neurons in the substantia nigra pars compacta and by the misfolding and aggregation of α-synuclein into cytoplasmic inclusions known as Lewy bodies, spreading across distinct brain regions in a temporospatial manner^5–9^.

Emerging models suggest that PD may originate through distinct pathological routes, with “body-first” and “brain-first” subtypes reflecting peripheral or central initiation of neurodegeneration^10–12^. Converging evidence implicates immune dysfunction and chronic inflammation as major contributors to disease onset and progression. Peripheral immune dysregulation and gut inflammation have been identified as risk factors for PD, and shared molecular mechanisms have been observed between PD and autoimmune conditions, particularly inflammatory bowel disease^13,14^. The gut–brain axis has therefore gained prominence as a bidirectional conduit linking peripheral immune responses to central neurodegeneration^15^.

Post-mortem analyses of PD brains reveal robust microglial activation, accompanied by upregulation of major histocompatibility complex class II (MHC-II), CD68, and toll-like receptors (TLRs)^16,17^. These markers reflect both resident microglial reactivity and the presence of infiltrating immune cells, including natural killer cells, monocytes, and neutrophils, as indicated by transcriptomic profiling of peripheral blood from PD patients^18^. Disruption of the blood–brain barrier (BBB) likely facilitates this infiltration, further amplifying local immune responses within the central nervous system (CNS)^19–21^. Sustained neuroinflammation in PD is evidenced by increased concentrations of cytokines and chemokines in the brain^22,23^ and cerebrospinal fluid^24^. The origin of these inflammatory mediators is complex and may involve both neural and peripheral sources. Extracellular vesicles (EV) have recently emerged as key mediators of intercellular communication and inflammatory propagation in neurodegenerative diseases. These membrane-bound nanoparticles, secreted by almost all cell types^25^, transport bioactive cargos, including misfolded proteins, nucleic acids, and lipids, that influence recipient cell function and gene expression^26^. In PD, EV have been implicated in the intercellular transfer of α-synuclein and in the modulation of glial activation and inflammatory signaling ^27–30^. Importantly, EV can traverse the BBB bidirectionally, providing a mechanistic link between peripheral and central immune processes^31^. Further, our group showed that distinctive pools of EV surface markers related to inflammatory and immune cells stratified patients with PD and multiples system atrophy according to the clinical diagnosis^32^. Despite these findings, the functional effects of plasmatic EV from PD patients on neuronal and microglial cells are not yet understood. Moreover, it is uncertain whether circulating EV can directly alter neuronal survival and modulate microglial activity, thereby linking peripheral immune dysfunction to neurodegeneration in PD.

Given that circulating EV largely originates from immune and endothelial cells, we hypothesized that plasma-derived EV from PD patients reflect peripheral immune activation and contribute to neuroinflammatory processes within the brain. Here, we examined the functional effects of patients-derived plasmatic EV in neuronal and glial cell cultures, assessing their capacity to induce direct neurotoxicity and to modulate microglia-neuronal interaction.

## Methods and Materials

### Patients’ recruitment and blood collection

Blood samples were taken from PD patients and sex-/age-matched healthy controls (HCs) participating in the NSIPD001 study. PD patients were prospectively recruited from the movement disorders outpatient clinic at Neurocenter of Southern Switzerland in Lugano; HCs were recruited among patient’s partners. The inclusion criteria for PD were (1) a definite clinical diagnosis according to the UK Parkinson’s Disease Society Brain Bank criteria^33^ and (2) >5 years since the clinical diagnosis. Exclusion criteria were significant comorbidities: diabetes, renal failure, thyroid pathology, vitamin B12 deficiency, HIV infection, syphilis, coagulopathy, fever, acute or chronic inflammatory diseases, and tumors. Ten mL of blood was collected into anticoagulant ethylenediamine tetraacetic acid (EDTA) tubes in the morning after a 4-hour fast.

### Plasma preparation

Fresh whole blood was centrifuged for 15 minutes at 1,600g at 10°C to eliminate cellular components. Cellular debris and platelets were removed by centrifuging the supernatant for 15 minutes at 3000g at 4 °C followed by 10,000g for 15 minutes and 20,000g for 30 minutes at 4°C, allowing the removal of larger EV and apoptotic bodies^32,34^. The obtained plasma was aliquoted and stored at −80°C. The storage period varied among samples according to the consecutive enrollment of subjects in the study, between July 2015 and January 2019 **(Figure 1a)**.

**Figure 1:**
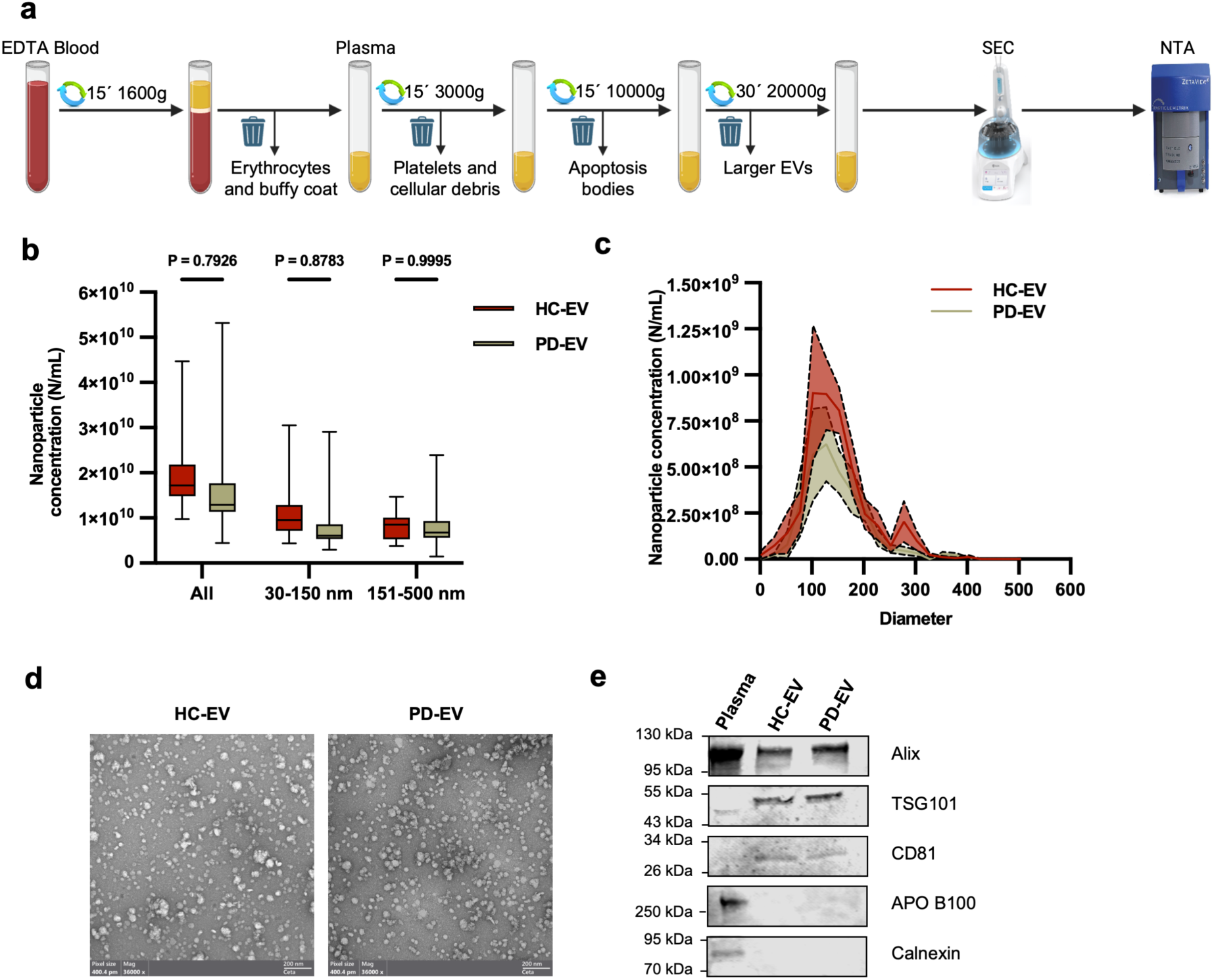
Characterization of plasmatic extracellular vesicles. **(a)** Schematic experimental setup for EV enrichment from plasma (created with BioRender). EDTA blood underwent serial centrifugation to eliminate cellular components and larger EVs. **(b)** PD-EV and HC-EV size profile distribution and nanoparticle concentration (N/mL) were quantified using NTA (smaller nanoparticles, 30-150 nm; larger nanoparticles, 151-500 nm) (n=14). **(c)** NTA representation of plasmatic EV from PD and HC. **(d)** TEM images of negatively stained EV from HC (left) and PD (right). **(e)** Western blot of EV showing EV-specific markers (CD81, Alix, and TSG101) and absence of contaminants (APO B100 and Calnexin).

### EV isolation

EV were isolated from human plasma using size-exclusion chromatography (SEC). Plasma samples (25 µL) from 10 PD and 10 age-/sex-matched HC were pooled for the analysis. The pooled plasma samples (250 µL) from PD patients and HC were diluted with 250 µL phosphate-buffered saline (PBS) and loaded onto the column (qEVoriginal columns 70nm, IZON) after washing with PBS. 30 × 0.5 mL fractions were collected, and. fractions 7-9 with higher concentration of EV and lower concentration of protein contaminants were used for the subsequent characterization and analyses.

### Nanoparticle tracking analysis (NTA)

The concentration (Nanoparticles/mL) and size distribution (nm) profiles of the EV were measured using Zeta View (Particle Metrix), equipped with a 40mW laser at 520nm and a CMOS camera. One µL of isolated plasma-EV was diluted 1:500 in PBS to obtain approximately 300 particles/frame. For each measurement, scatter mode was performed at a cell temperature of 25 °C and videos of 11 cell positions were acquired with the following parameters: sensitivity, 80; shutter, 100; Minimum Brightness, 30; Maximum Area, 1000; and Minimum Area, 10. All parameters were held constant across samples to minimize variability. Videos were analyzed using the built-in ZetaView 8.02.31 software with an embedded laser of 40 mW at 520 nm and a CMOS camera. The number of completed tracks in NTA measurements was always greater than the proposed minimum of 1000, which helps minimize data skewing due to a single large particle.

### Transmission electron microscopy (TEM)

EV morphology was evaluated by negative staining TEM. Briefly, 2 µL of the EV samples were absorbed onto a glow-discharged 300-mesh carbon film grid. After 5 minutes, the excess sample was removed, and the grid was negatively stained with 2% Uranyl-acetate diluted in 1:4 in ddH_2_O. After 2 minutes, the UA was removed, and the grid was air-dried for 30 minutes. EV were examined by a Talos L120C (Thermo Fisher Scientific) operating at 120 kV. Images were acquired using a 4 K × 4 K Ceta CMOS camera (Thermo Fisher Scientific).

### Cell culture material and reagents

Human neuroblastoma SH-SY5Y cells (Sigma, 94939304) were grown at 37 °C in a humidified atmosphere, 5% CO_2_ in Dulbecco’s Modified Eagle Medium (DMEM, Thermo Fisher Scientific) supplemented with 10% fetal bovine serum (FBS, Sigma), 1% penicillin-streptomycin (P/S, Thermo Fisher Scientific), 1X GlutaMAX (Thermo Fisher Scientific), and 1% non-essential amino acids (NEAA, Thermo Fisher Scientific). Human microglia HMC3 cells (ATCC, Cat. No. CRL-3304) were maintained at 37 °C, 5% CO_2_ in a DMEM/F-12 (Thermo Fisher Scientific) supplemented with GlutaMAX, 10% v/v FBS, 1% NEAA, and 1% P/S. All cell lines were routinely tested for the presence of mycoplasma.

### Western blot analysis

HMC3 cells and EV were lysed in 1X RIPA buffer (Sigma) containing 1X protease and phosphatase inhibitors. A QuantiPro^TM^ BCA assay kit (SIGMA) was used to calculate protein concentration. In total, 20 µg of proteins were heated at 95 °C for 10 minutes with SDS loading buffer (6X) and separated on 4-20% Mini-PROTEAN® TGX^TM^ Precast Gel (BioRad) and transferred onto a PVDF membrane using Trans-Blot Turbo (BioRad) for 25 minutes. Membranes were blocked using Intercept (TBS) blocking buffer for 1 hour and then incubated overnight at 4°C with the following primary antibodies: anti-CD81 (1:500, Abcam, ab109201), anti-TSG101 (1:1000, Abcam, ab83), anti-ApoB (1:500, Abcam, ab20737), anti-Calnexin (1:500, Abcam, ab133615), anti-Alix (1:500, Abcam, ab186429), anti-p16 (1:1000, Proteintech, 10883-1-AP), anti-p21 (1:1000, Proteintech, 10355-1-AP), and anti-β-Actin (1:1000, SIGMA, A1978), overnight at 4°C. The following day, after three 10-minute washing steps in Tris-buffered saline with Tween-20 (TBS-T), the blots were incubated with secondary IRDye antibodies for 1 hour at room temperature (RT). After three 10-minute washes in TBS-T, blots were visualized and analyzed using LICOR Odyssey CLx imaging system (LI-COR Biosciences).

### Differentiation of SH-SY5Y cells

SH-SY5Y neuroblastoma cells were differentiated into neuron-like cells using methods described in previous studies^35–38^. Briefly, undifferentiated SH-SY5Y, when reached 70-80% confluence, were trypsinized with TrypLE Express enzyme (Thermo Fisher Scientific) and seeded in DMEM/F-12 supplemented with GlutaMAX, 10% v/v FBS, 1% NEAA, and 1% P/S at a density of 2.5 × 10^4^ cells/well in 12-well plates previously coated with poly-D-lysine (Sigma) and laminin (Sigma). Twenty-four hours after plating, the complete medium was replaced with a reduced medium (1% FBS) containing 10 µM retinoic acid (ATRA) to promote differentiation and the neuronal phenotype. The cells were allowed to grow in 10 µM ATRA-reduced medium for at least 5 days, with the medium refreshed every other day. 50 ng/mL BDNF was added for an additional 3-5 days after ATRA treatment to enhance the effect of ATRA on neuronal differentiation. The SH-SY5Y cells were terminally differentiated into neuron-like cells on day 10 and used for further studies.

### EV treatment on SH-SY5Y neurons

SH-SY5Y neurons were treated with EV from PD, HC or PBS at a final concentration of 6.5 χ 10^8^ EV/mL and 5 χ 10^8^ EV/mL, and readouts were performed at 24 hours.

### EV treatment on HMC3 cells

HMC3 cells were first synchronized overnight in serum-free medium and then stimulated with PD-EV, HC-EV, or PBS at a final concentration of 5 χ 10^8^ EV/mL under serum-free medium for 48 and 96 hours. Proinflammatory and senescence-specific phenotypes were evaluated. Conditioned medium (CM) was isolated and centrifuged at 3000g for 10 minutes to remove dead cells and debris, and in a subset of experiments followed by ultrafiltration with a 30kDa filter at 3000g for 30 minutes to remove EV. SH-SY5Y neurons were then treated with CM or EV-depleted CM for 24 hours.

### Uptake of EV by flow cytometry

SH-SY5Y cells were seeded in 12-well plates at a density of 2.5 × 10^4^ cells/well for differentiation. Following differentiation, cells were washed with PBS and co-cultured with PD-EV, HC-EV, or PBS, previously labeled with DiR cell-labeling dye. SH-SY5Y neurons were harvested with TrypLE Express (Thermo Fisher Scientific) and centrifuged for 5 minutes at 350g. Pellets were resuspended in 200 µL PBS with 4% FBS. Cells were acquired by flow cytometry (750 nm, CytoFLEX) at 6, 12, and 24 hours, and analyzed with FlowJo 10.9.0.

### Immunofluorescence staining

After treatment, cells were fixed with 4% paraformaldehyde for 15 minutes, permeabilized, and blocked with 5% Normal Goat Serum (NGS) and 0.3% Triton X-100 in PBS for 30 minutes. Cells were stained with primary antibody (diluted in 0.5% NGS and 0.3% Triton X-100 in PBS) for 1 hour at 37°C, thoroughly washed with PBS, and stained with the secondary antibody (diluted in 0.5% NGS and 0.3% Triton X-100 in PBS) for 1 hour at RT, and followed by nuclear staining with DAPI (1:5000) for 5 minutes at RT. The following primary antibodies were used: rabbit antibody anti-beta-tubulin III (TUBB3, 1:1000, ab18207, Abcam), rabbit antibody anti-IBA1 (1:250, PA5-27436, Thermo Fisher Scientific), and mouse antibody anti-alpha-synuclein (Syn211, 1:500, ab80627, Abcam). Stained cells were imaged using a fluorescence microscope (Nikon ECLIPSE Ti2).

### Neurite length calculation and morphology analysis

Neurite length of SH-SY5Y neurons (using the NeuronJ plugin) and morphological analysis of the human microglial cell line HMC3 were performed using ImageJ software ^39,40^. Four to five regions of interest images were randomly selected from each well. Neurites were manually traced, and their lengths (μm) were measured. SH-SY5Y neurons were considered to be differentiated if they met the following morphological criteria: the neurite length should exceed 15 µm, and the neurite should extend its outgrowth at least two-fold higher than the soma diameter **(Figure S2b)**. Activation of HMC3 cells was characterized by their morphology, as determined by the microglial lineage (IBA1) marker and cell surface area. Images were analyzed by a rater blinded to experimental conditions.

### Annexin V/PI apoptosis Assay

After treatment, cells were harvested using TrypLE Express and centrifuged at 1200 rpm for 5 minutes. Cell pellets were resuspended in 1X 100µL assay buffer. Per reaction, 1µL Annexin V-FITC (BD Biosciences, cat. no. 556547) and 1µL Propidium Iodide (BD Biosciences, cat. no. 556547) were incubated for 15 minutes at RT in the dark. Samples were analyzed using flow cytometry (CytoFLEX S).

### Cytokine Expression Assay

The Human Neuro Discovery Array C2 was used to investigate the cytokine release from EV-treated HMC3 cells. In brief, HMC3 cells were grown in a 12-well plate. After 24 hours, cells were treated with PD-EV and HC-EV at a final concentration of 5 χ 10^8^ EV/mL in a reduced EV-depleted serum medium to avoid lipoprotein contamination. After 24 hours, CM was isolated and centrifuged at 3000g for 30 minutes at 10°C to remove particulates. Cytokine array membranes precoated with cytokine antibodies were blocked using a blocking buffer for 30 minutes at RT^41,42^. The blocking solution was replaced with CM isolated from EV-treated HMC3 cells and incubated for 5 hours at RT. Membranes were washed multiple times and incubated with a biotinylated antibody cocktail for 2 hours at RT. Fluorophore-conjugated streptavidin was added and incubated for 2 hours at RT, after which the membranes were visualized using the LI-COR machine. Cytokine array results were analyzed using the manufacturer’s analysis tool.

### Real-time PCR

Total RNA was extracted from HMC3 cells using 1 mL TRI-Reagent (Sigma-Aldrich) according to the manufacturer’s instructions. 150-500ng of RNA were reverse transcribed with the help of GoScript Reverse Transcription system kit (Promega) according to the manufacturer’s instructions. Real-time PCR was performed with the following mix: 5 µl SsoAdvanced universal SYBR green supermix 2x (BioRad), 3 µl DEPC water, 1 µl cDNA diluted 1:5 in DEPC water, 0.5 µl forward primers (10 µM) and 0.5 µl reverse primers (10 µM). Amplification and detection of specific genes were performed in triplicate using the QuantStudio 1 Real-Time PCR system (Applied Biosystems^TM^). The cycle threshold (Ct) for each gene was normalized to the geometric mean of the housekeeping genes RPL27 and GAPDH (ΔCt). To compare relative gene expression levels among HC- and PD-EV treatments, ΔΔCt values in EV-treated HMC3 cells were calculated as the difference between the ΔCt value in these groups and the ΔCt value in PBS-treated HMC3^43^. The primers used in this paper are as follows: *p16^INK^*^4a^ 5^’^-CTTCGGCTGACTGGCTGG-3^’^ (forward) and 5^’^-TCATCATGACCTGGATCGGC-3^’^ (reverse); and *p21^WAF^*^1^*^/CIP^*^1^ 5^’^-CACCTCACCTGCTCTGCTGC-3^’^ (forward) and 5^’^-GCTGGTCTGCCGCCGTTTT-3^’^ (reverse); *SERPINE1* 5^’^-TTGCAGGATGGAACTACGGG -3^’^(forward) and 5^’^-GTGGCAGGCAGTACAAGAGT -3^’^(reverse); *GAPDH* 5^’^-TGCACCACCAACTGCTTAGC-3^’^(forward) and 5^’^-GGCATGGACTGTGGTCATGAG- 3^’^(reverse); *RPL27* 5^’^-TGGTAGGGCCGGGTGGTTG -3^’^(forward) and 5^’^-ACTTTGCGGGGGTAGCGGTC-3^’^(reverse).

### Liquid chromatography-tandem mass spectrometry (LC**-**MS/MS)

#### Protein extraction and enzymatic digestion

EV lysis and protein extraction were performed in 4% SDS in 100 mM Tris pH 7.6 with 10 mM dithiothreitol by sonication at 4°C in a Bioruptor (Diagenode, 15 cycles, 30s on, 30s off, high mode) and incubation at 95°C for 10 minutes. Proteins were alkylated with 50 mM iodoacetamide for 30 minutes at RT and then precipitated overnight in 80% cold acetone. The next day, proteins were pelleted by centrifugation at 13,000 rpm for 20 minutes at 4°C, washed twice with 80% cold acetone, and dried at 40°C. Protein pellets were resuspended in 8M urea in 50 mM ammonium bicarbonate (ABC) through Bioruptor sonication, and digestion was carried out with LysC (Wako Fujifilm, 1:100 w/w) for 2 hours at RT. The digestion buffer was then diluted to 2M urea with 50 mM ABC, and trypsin (Thermo Fisher Scientific, 1:100 w/w) was added for overnight digestion at RT. Digestion was halted by adding acetonitrile (ACN) to 2% and trifluoroacetic acid (TFA) to 0.3%. The samples were then cleared by centrifugation for 5 minutes at maximum speed. Peptides were purified on C18 StageTips^44^, and eluted with 80% ACN, 0.5% acetic acid. Finally, the elution buffer was evaporated by vacuum centrifugation, and the purified peptides were resuspended in 2% ACN, 0.5% acetic acid, and 0.1% TFA for single-shot LC-MS/MS measurements.

#### LC-MS/MS analysis

Samples were analyzed using a nanoElute2 HPLC system (Bruker) coupled via a nanoelectrospray source (Captive Spray Source, Bruker) to a timsTOF HT mass spectrometer (Bruker). Peptides were loaded in water/0.1% formic acid into a 75 µm inner diameter, 25 cm long column in-house packed with ReproSil-Pur C18-AQ 1.9 µm resin (Dr. Maisch HPLC GmbH) kept at 50°C in a column oven, and eluted over a 60-min linear gradient between 2 and 35% ACN/0.1% formic acid at a flow rate of 300 nl/minutes. The mass spectrometer was operated in a data-dependent (DDA)-PASEF moFNde, with 10 PASEF ramps, each with an accumulation and ramp time of 166 ms, covering a 100-1700 m/z range and a 0.60-1.60 Vs/cm² mobility range, for a total cycle time of 1.89 s. Collision energy was ramped linearly from 20 eV at 0.60 Vs/cm^2^ to 59 eV at 1.60 Vs/cm^2^. The signal intensity threshold was set at 1,000, and precursors were actively excluded for 0.4 min after reaching a target intensity of 20,000. Precursors with a charge of up to 5 were selected for fragmentation using Bruker’s default active precursor region filter. The high sensitivity detection mode for low sample amounts was enabled.MS raw files were processed using MaxQuant software version 2.4.2.0^45^. The Andromeda search engine^46^ was employed to identify peptides and proteins with a false discovery rate of < 1% from the Human UniProt database (February 2024) and a common contaminants database (247 entries). Enzyme specificity was set as “Trypsin/P” with a maximum of 2 missed cleavages and a minimum length of 7 amino acids. N-terminal protein acetylation and methionine oxidation were set as variable modifications, and cysteine carbamidomethylation as a fixed modification. Label-free protein quantification (LFQ) was calculated by the MaxLFQ algorithm^47^ with a minimum peptide ratio count of 1, and absolute protein abundances were calculated with the iBAQ method. The proteinGroups.txt table was pre-processed by removing proteins identified only by site, reverse hits, and potential contaminants. Both iBAQ and LFQ intensities were log2-transformed.

### Functional pathway enrichment analysis

Functional pathway enrichment was conducted using the methodology outlined by Reimand et al^48,49^. We generated a generic enrichment map file (GEM) for pathway analysis using g:Profiler, which conducts functional profiling of proteins from a cytokine expression assay and proteomic analysis of EV (https://biit.cs.ut.ee/gprofiler/)^50^. In addition to Gene Ontology (biological process, molecular function, and cellular component), pathways from the Reactome databases were also included. The results were visualized as an enrichment bubble plot generated with ggplot2 in R, where the bubble color represents the -log10 (adjusted p-value) and the size represents the number of enriched proteins.

### Statistical analysis

All data were analyzed using PRISM analysis software (GraphPad Prism 10). Variable distribution was assessed using the Kolmogorov-Smirnov and Shapiro-Wilk tests. Normally distributed variables are represented as mean ± standard error of the mean and analyzed using Student’s two-tailed t-test or one-way ANOVA with Tukey’s post hoc test, multiple comparisons, or two-way ANOVA with Bonferroni multiple comparisons (for multiple groups). Non-normally distributed variables are represented as median with interquartile range and were analyzed using the Mann-Whitney U-test (two groups) or Kruskal-Wallis test with Dunn’s multiple comparison correction (multiple groups). P<0.05 were considered significant.

## Results

### Characterization of EV

EV isolated from PD and HC plasma were characterized according to the MISEV guidelines^25^. No differences were observed between the two groups in concentration, expressed as nanoparticles/mL **(Figure 1b),** and mean diameter by NTA (160.21±3.37 nm in PD and 155.07± 4.63 nm in HC) (**Figure S1**). The size distribution curves for HC-EV and PD-EV were similar **(Figure 1c)**. TEM images revealed a spherical shape characteristic of EV, with a size below 200 nm, corresponding to small EV (**Figure 1d**). Western blot analysis showed the absence of contaminants (APOB and calnexin) and the presence of EV-specific luminal markers (TSG101 and Alix) and EV-specific tetraspanin (CD81) in the isolated EV **(Figure 1e and S5a)**.

### Differentiation of Neuroblastoma SH-SY5Y into mature neurons

Neuroblastoma SH-SY5Y cells were treated with ATRA and brain-derived neurotrophic factors (BDNF) to induce differentiation into mature neurons. Differentiation was analyzed using the neuronal marker beta-tubulin III staining on day 10. Undifferentiated SH-SY5Y cells exhibited a dense, flat, epithelial-like morphology with few small outward processes. In contrast, the differentiated neuron-like cells displayed extensive, interconnected, and elongated neurite projections (Figure S2a-c). Additionally, alpha-synuclein was detected in the axons of the differentiated SH-SY5Y neurons (**Figure S2a).**

### EV were internalized by SH-SY5Y

First, we investigated the internalization of PD-EV and HC-EV. SH-SY5Y neurons were incubated with DiR-labeled PD-EV, HC-EV, and PBS (negative control), and their internalization was quantified by flow cytometry **(Figure 2)**. After 6 hours, SH-SY5Y cells treated with PD-EV and HC-EV showed increased DiR fluorescence, whereas PBS-treated SH-SY5Y neurons exhibited no signal. We estimated that 30% of SH-SY5Y neurons internalized EV after 6 hours, with up to 50% after 24 hours. EV internalization increased in a time-dependent manner, with higher levels at 24 hours. Notably, no differences were found in internalization between PD-EV and HC-EV at any time point.

**Figure 2:**
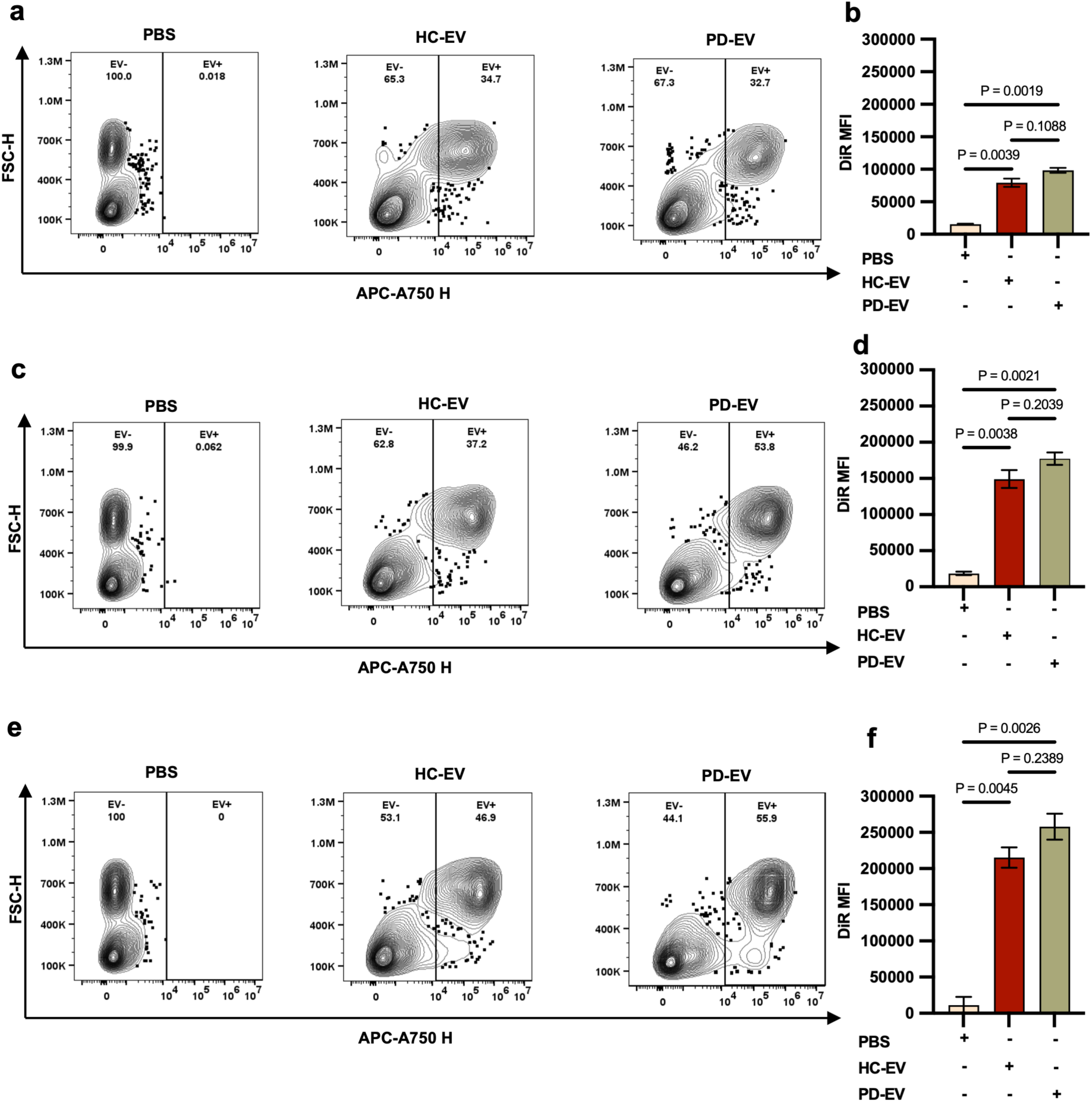
Uptake analysis of plasma-derived EV. Internalization of DiR-labelled HC-EV, PD-EV, and PBS by SH-SY5Y neurons at 6 hours **(a),** 12 hours **(c)**, and 24 hours **(e)** by flow cytometry. **(b, d, f)** Bar graph showing median fluorescence intensity of DiR-labelled EV internalized by SH-SY5Y neurons using FlowJo software. Data are represented as mean ± SEM. Results correspond to 2-3 replicates from at least two independent differentiations, and P-values are shown in the figures.

### PD-EV reduced neurite length in SH-SY5Y

SH-SY5Y neurons were incubated with PD-EV and HC-EV for 24 hours at 5 × 10^8^ EV/mL and 6.5× 10^6^ EV/mL (**Figure 3a and Figure S3a**). SH-SY5Y neurons incubated with higher dose of PD-EV showed a significant reduction in neurite length, whereas neurons incubated with HC-EV and PBS showed no detectable effect (**Figure 3b-c and Figure S3a)**. Since no notable difference was found in neurite length reduction between the two concentrations of EV (6.5 × 10^6^ and 5×10^8^ EV/mL) **(Figure S3b-c)**, the highest, closer to the in vivo plasmatic EV concentration (∼10^10^)^51^, was chosen for further downstream analysis.

**Figure 3:**
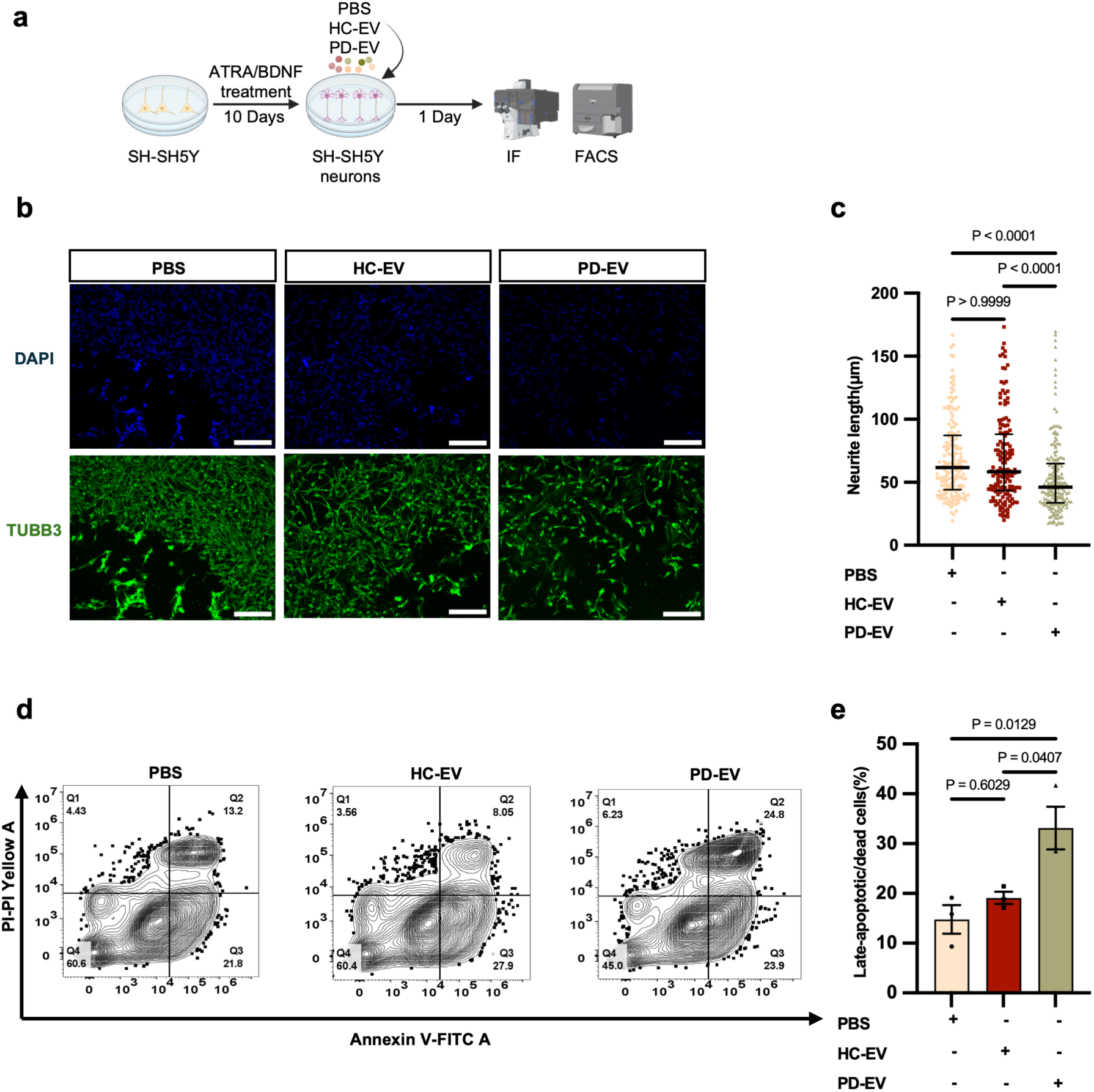
Analysis of neurite length reduction and cell death in SH-SY5Y neurons after treatment with EV. **(a)** Schematic experimental setup of EV treatment on SH-SY5Y neurons (created with BioRender). **(b)** Immunostaining of SH-SY5Y neurons with DAPI (blue) and TUBB3 (green) after EV treatment. Scale bar: 100 µm. **(c)** Quantification of neurite length of SH-SY5Y neurons after 24 hours of treatment with PD-EV, HC-EV, and PBS (n=150-200 neurons per group). The graph presents all the data points, with the median represented as a straight line and the IQR as an error bar for neurite length **(d)** Flow cytometry evaluation of SH-SY5Y neurons after 24 hours of EV treatment using Annexin V FITC (x-axis) and propidium iodide (y-axis). Q1: dead cells (PI positive), Q2: late-stage apoptotic cells (Annexin V and PI positive), Q3: early-stage apoptotic cells (Annexin V positive), Q4: live cells (Annexin V and PI negative). Results correspond to 3 replicates from at least three independent differentiations. (e) Bar graph showing percentage of cells in Q1 and Q2 stages after 24 hours treatment with EV. Data are represented as mean ± SEM for apoptosis. P-values are shown in the figures.

After EV treatment, we performed Annexin V/PI staining followed by flow cytometry to identify apoptotic and dead cells. As neuronal differentiation is intrinsically a stressful process, we expected to observe at least one apoptotic marker under untreated conditions during differentiation, which aligns with previous studies^52,53^. However, following PD-EV treatment, we observed a significant increase in the populations of both late-stage apoptotic (annexin V+/PI+) and dead (annexin V-/PI+) cells. On average, 33.41% of SH-SY5Y neurons were in late-stage apoptosis/dead at 24 hours **(Figure 3d-e)**.

### PD-EV induced an activated state in human microglia HMC3

Different phenotypes of microglia are generally classified based on their morphological features. “Resting” microglia are highly ramified with small cell bodies; whereas upon activation, microglia acquire a more amoeboid-like shape with larger sizes and bigger cell bodies^54^. HC-EV- and PBS-HMC3 cells displayed the typical morphology of resting microglia, with higher bipolar ramification and smaller cell bodies **(Figure 4a)**. Upon treatment with PD-EV, HMC3 cells transformed into an amoeboid-like phenotype with larger cell bodies, fewer ramifications, and higher surface area compared to PBS and HC-EV-treated HMC3 cells **(Figure 4b-c)**.

**Figure 4:**
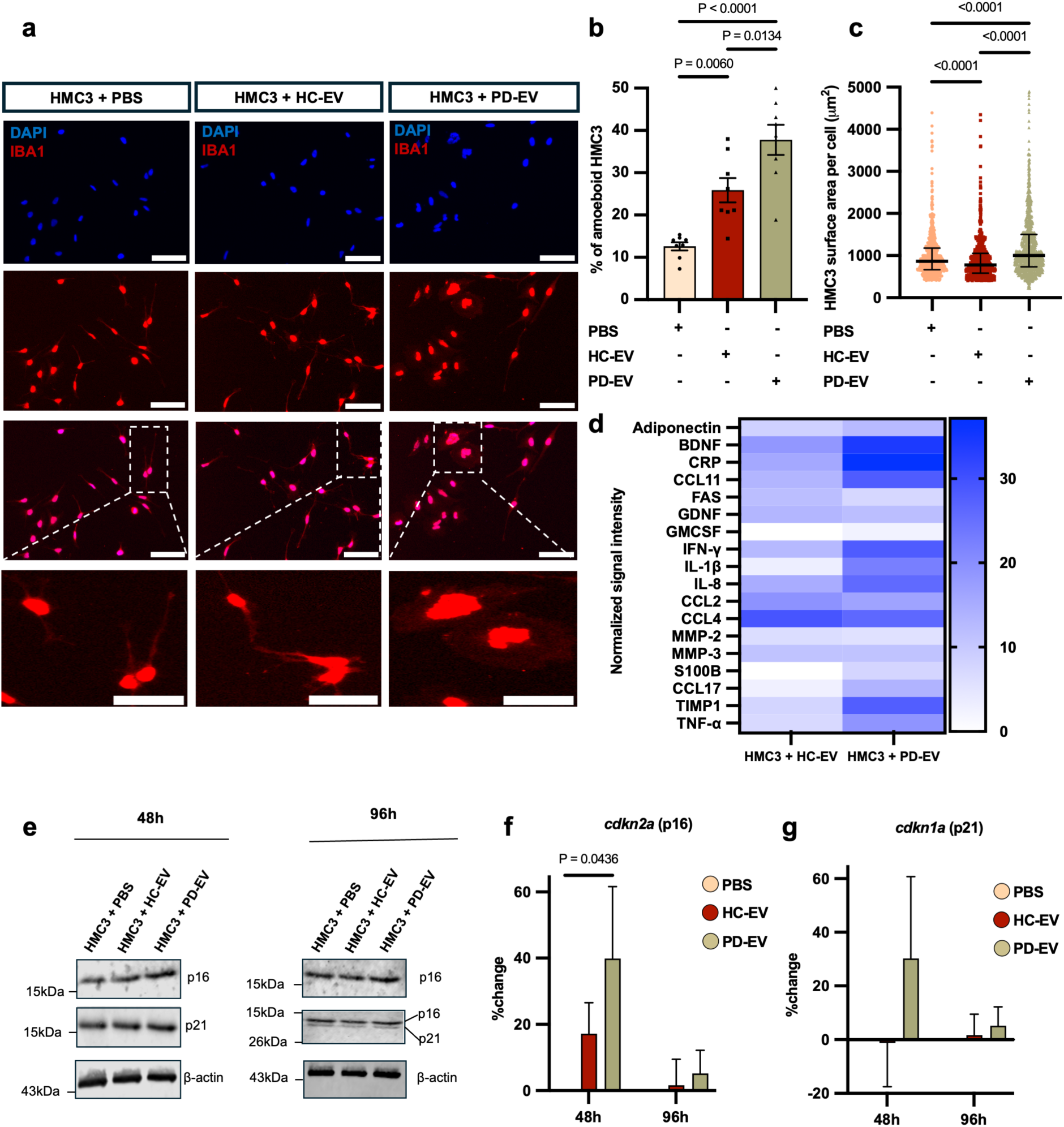
Morphological, cytokine expression and senescence-associated markers analysis of HMC3 cells after EV treatment. **(a)** Immunostaining of HMC3 cells with DAPI (blue) and IBA1 (red) after 24 hours of EV treatment. Scale bar: 100 μm; zoom scale bar: 50 μm **(b-c).** Percentage and surface area of activated HMC3 cells after 24 hours of EV treatment. Data are represented as mean ± SEM and median with interquartile range (n=8 images per group). P-values are represented in the figures. **(d)** Heatmap showing normalized signal intensity of cytokine expression in HMC3 cells treated with PD-EV- and HC-EV**. (e)** Western blot quantification of p16^INK4a^ and p21^WAF1/CIP1^ in HMC3 cells after 48 and 96 hours of EV treatment (n=3). (f-g) Semi-quantification of p16^INK4a^ and p21^WAF^^1/CIP1^ expressed as percentage change: (Normalized signal (EV-treated))/ (Normalized signal (PBS control)-1) *100. Data are represented as mean ± SEM. P-value is represented in the figures.

### PD-EV induced proinflammatory cytokines and senescence-like phenotypes in HMC3

CM from HMC3 cells treated with PD-EV and HC-EV were analyzed by cytokine array. Proinflammatory senescence-associated secretory phenotypes (SASP), including TNF-α, IL-1β, IFN-γ, IL-8, and CCL11^55^, were elevated in PD-EV-HMC3 compared to those in the HC-EV-HMC3. Chemokines and regulatory factors such as CCL11, CCL17, TIMP1, and GM-CSF, which are involved in aging-related inflammation and alternative immune activation, were also elevated in PD-EV-HMC3 cells; BDNF, adiponectin, and CRP were increased after PD-EV treatment. In contrast, FAS levels were higher in HC-EV-HMC3 cells **(Figure 4d)**. Of interest, S100B resulted increased after PD-EV treatment. While astrocytes are the primary source of S100B in the brain, microglia are both producers and major targets of this protein, playing a role in its regulation during inflammation and injury^56^.

Given the upregulation of the SASP, we investigated other senescence-related features in HMC3 cells. HMC3 cells treated with PD-EV showed a significant increase in p16^INK4a^ protein levels at day 2, which declined over time at day 4. Additionally, p21^WAF1/CIP1^ showed an upward trend at day 2, albeit not significant (**Figure 4e-g and S5b-c**). RT-PCR performed on HMC3 cells treated with PD-EV vs HC-EV further showed an upward trend of p16^INK4a^ and SERPINE1 expression after PD-EV treatment at both day 2 and 4, with a slight increase of p21^WAF1/CIP1^ at day 4 (**Figure S4a-c**).

To assess the functional pathways associated with upregulated cytokines in the CM of PD-EV-HMC3 cells, we conducted a functional pathway enrichment analysis using g:Profiler, which was curated from the Gene Ontology and Reactome databases. These cytokines were associated with both metabolic processes and immune system, in particular with IL-17 signaling pathway, cytokine-cytokine receptor interaction, type I diabetes mellitus, non-alcoholic fatty liver disease, inflammatory bowel disease, NF-κB signaling pathway, and Toll-like receptor signaling pathway **(Figure 5a-b)**.

**Figure 5:**
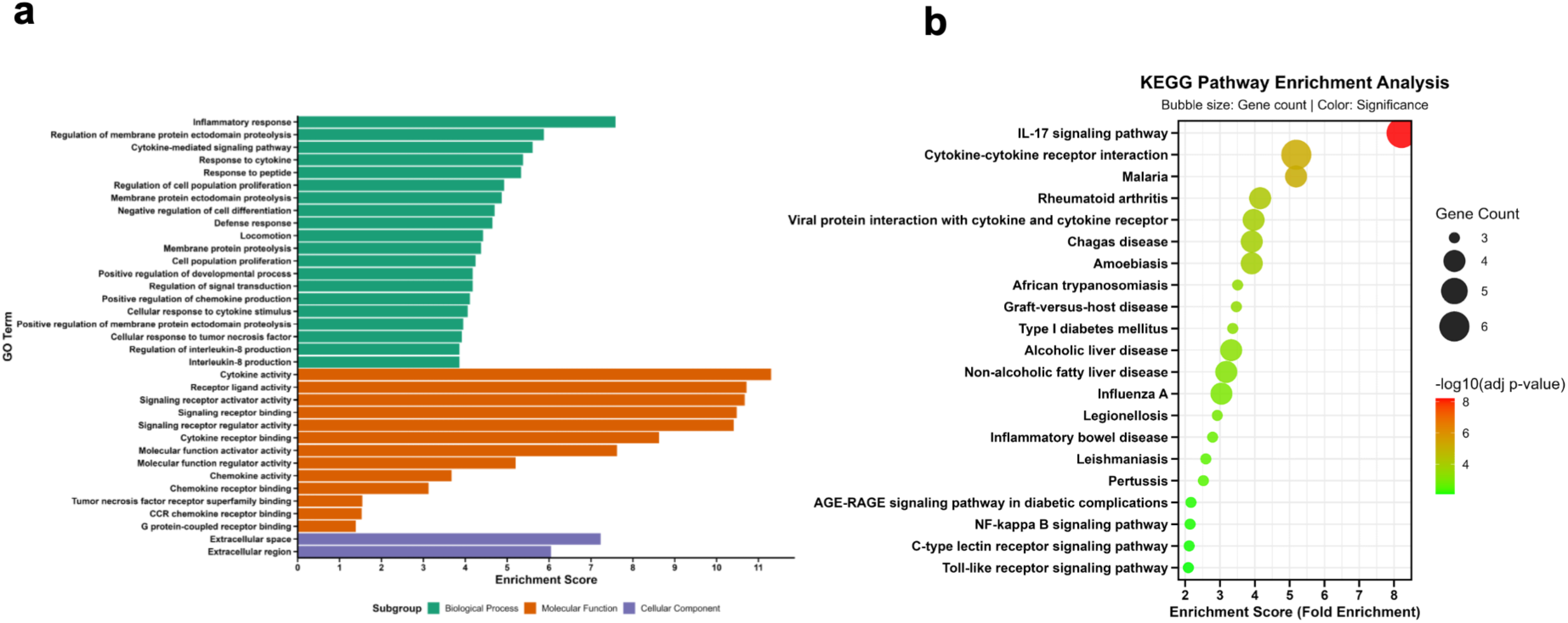
Functional pathways of HMC3 cells after EV treatment. **(a)** GO (biological process, molecular function and cellular component) analysis of enriched cytokines. X-axis represents enrichment score. **(b)** Signaling pathway enrichment analysis of cytokines by KEGG database in g: Profiler.

### Conditioned media from PD-EV-treated HMC3 induced apoptosis in SH-SY5Y

To study the effect of activated microglia on neuronal cells, we treated SH-SY5Y neurons with HMC3 CM that had been previously incubated with PD-EV, HC-EV, or PBS (**Figure 6a**). To account for possible residual EV from treatment, we also evaluated the effects of EV-depleted CM (EV^-^ CM) on SH-SY5Y neurons. A significant reduction in neurite length (**Figure 6b-c**) and an increase in late-stage apoptosis/dead (**Figure 6d-e**) were observed with CM from PD-EV-HMC3 cells compared to other conditions. In addition, SH-SY5Y neurons exposed to EV^-^ CM from PD-EV-HMC3 also showed significantly reduced neurite length (**Figure 6f-g**) and increased proportion of cells in late-stage apoptosis/dead (**Figure 6h-i**), although levels were slightly lower than those observed with EV-containing CM. Notably, when comparing the direct effect of PD-EV on SH-SY5Y neurons with the indirect effect of CM or EV^-^ CM from PD-EV-HMC3 cells, a distinct toxicity pattern was observed. Direct PD-EV treatment resulted in a greater reduction in neurite length than CM or EV^-^CM **(Figure 6j)**. However, late-stage apoptosis was higher in SH-SY5Y neurons treated with CM (56.9%) and EV^-^ CM (51.9%) than in those treated with direct PD-EV (33.1%) (**Figure 6k**).

**Figure 6:**
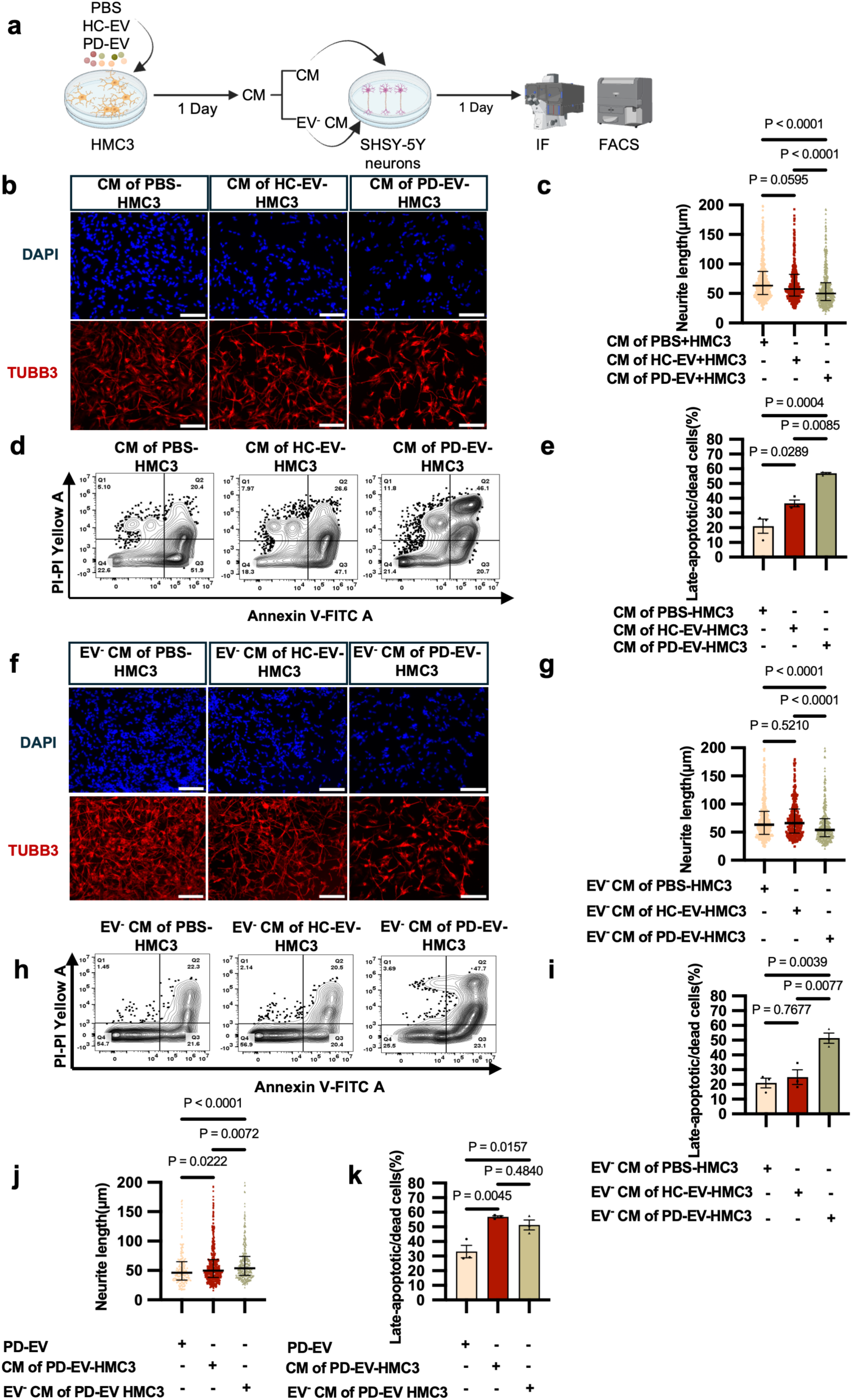
Neurite length reduction and cell death in SH-SY5Y neurons after treatment with CM and EV^-^ CM of PD-EV, HC-EV, and PBS-treated HMC3 cells. **(a)** Schematic experimental setup of EV treatment on HMC3 cells followed by CM isolation and SH-SY5Y neurons treatment. **(b,f)** Immunostaining with DAPI (blue) and TUBB3 (red) of SH-SY5Y neurons after treatment with CM and EV^-^ CM. Scale bar: 200 µm. **(c,g)** Neurite length quantification of SH-SY5Y neurons after 24 hours treatment with CM and EV^-^ CM (n= 550-580 neurons per group). **(d,h)** Flow cytometry analysis of SH-SY5Y neurons exposed to CM and EV^-^ CM for 24 hours. Annexin V FITC (x-axis) and propidium iodide (y-axis). Q1: dead cells (PI positive), Q2: late-stage apoptotic cells (Annexin V and PI positive), Q3: early-stage apoptotic cells (Annexin V positive), Q4: live cells (Annexin V and PI negative). **(e,i)** Bar graph showing percentage of cells in Q1 and Q2 stages after 24 hours treatment with CM and EV^-^CM. Results correspond to 3 replicates from at least three independent differentiations. **(j)** Neurite length and **(k)** apoptosis comparison of SH-SY5Y neurons treated with CM and EV^-^ CM, respectively. Scatter plots show all data points, with median as a straight line and the IQR as an error bar. Bar plots show mean ± SEM. P-values are represented in the figures.

### Proteomics landscape of PD-EV identified complement activation and immune response pathways

Proteomic analysis of pooled PD-EV and HC-EV identified 88 proteins, 20 of which were exclusively present in PD-EV (**Figure 7a**). PD-EV showed higher levels of complement proteins C1QC, C4, C5, and CFH; immunoglobulins IGKV3D-11, IGKC, IGHG1, IGHA, and IGKV4-1 IGLC; cytoskeleton regulators CFL1 and CALM; acute-phase proteins SERPINA1, HP, and A2M; extracellular matrix regulators VWF and THBS1; and stress response protein QSOX1. On the contrary, CLU, FBLN1, APOD, EPB42, and LGALS3BP were downregulated in PD-EV (**Figure 7b-c**). Detailed protein data are provided in the **supplementary data 1**. Focusing on the more abundant proteins, we conducted Gene Ontology analysis (Biological Process, Molecular Function, Cellular Component, and Reactome pathways). We found the highest enrichment scores in complement cascade (p= 9.8 E^-15^), innate immune system (p=1.85 E^-10^), binding and uptake of ligands by scavenger receptors (p=2.52 E^-7^), and of interest, post-translational protein phosphorylation (p=2.23 E^-05^) (**Figure 7d**). The biological process showed enrichment in complement activation (p=2.13 E^-13^), especially the classical pathway (p=3.09 E^-08^), humoral (p=1.63 E^-09^), and adaptive (p=9.03 E^-09^) immune responses, as well as coagulation (p=0.0273) (**Figure 7e**), suggesting its involvement in inflammation. Of note, cellular component analysis showed that proteins were enriched in blood microparticle (p=1.03 E^-23^), extracellular space (p=5.46 E^-21^), extracellular vesicle (p=1.09 E^-11^), and extracellular organelle (p=1.10 E^-11^), confirming EV enrichment and the quality of EV-purified fractions used in these experiments (**Figure 7f**). Molecular function analysis revealed enrichment in endopeptidase regulator activity (p=6.36 E^-07^), enzyme inhibitor activity (p=5.07 E^-06^), and complement binding (p=0.001397) (**Figure 7g**).

**Figure 7:**
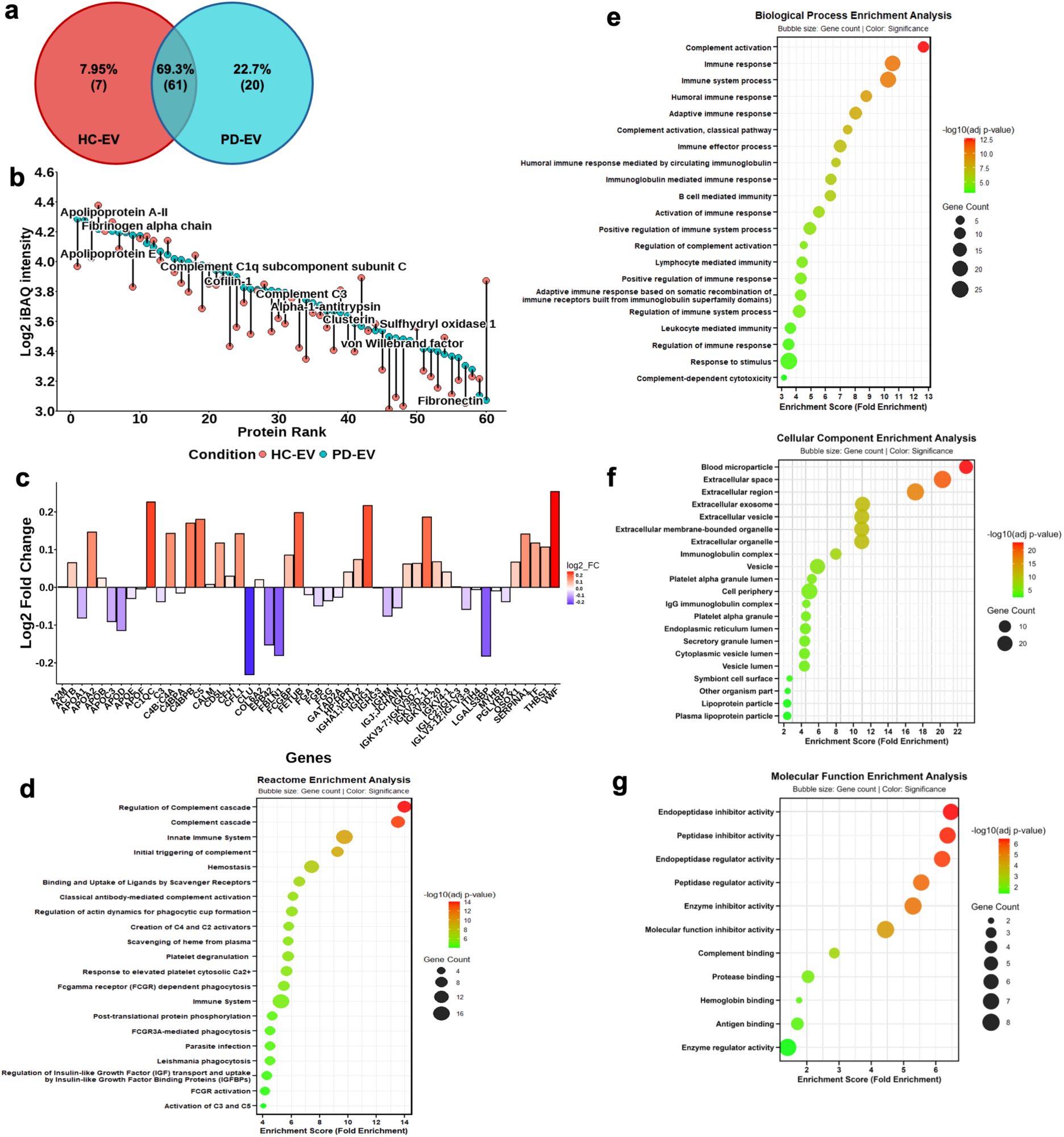
Proteomic profiling of EV. **(a)** Venn diagram of quantified proteins in HC-EV and PD-EV. **(b)** Protein rank plot showing proteins detected in PD- and HC-EV, ranked from highest to lowest log_2_ iBAQ intensity. Red represents proteins in HC-EV and green represents proteins in PD-EV. **(c)** Comparison of proteins log_2_ fold change between HC-EV and PD-EV. The x-axis represents genes and the y-axis log_2_ fold change. GO analysis of upregulated proteins in PD-EV vs HC-EV. **(d)** Reactome pathway. **(e)** Biological function. **(f)** Cellular component. **(g)** Molecular function. The x-axis represents enrichment score (-log_10_ of the adjusted p-value).

Conversely, proteins downregulated in PD-EV showed the highest enrichment scores in the pathways, such as platelet degranulation (p=2.04 E^-07^), response to elevated platelet cytosolic Ca^2+^ (p=2.69 E^-07^), plasma lipoprotein remodelling (p=5.58 E^-05^), and homeostasis (p=0.001555) (**Figure 8a**), suggesting their involvement in platelet and lipoprotein regulation. The biological processes for these proteins showed enrichment in blood coagulation and fibrin clot formation (p=1.96 E^-06^), reverse cholesterol transport (p=3.28 E^-06^), protein activation cascade (p=4.16 E^-06^), and amyloid-beta clearance (p=0.0096) (**Figure 8b**). The molecular function enrichment was in lipoprotein particle receptor binding (p=3.48 E^-06^), cholesterol binding (p=4.23 E^-05^), Phosphatidylcholine-sterol O-acyltransferase activator activity (p=0.003338), and amyloid-beta binding (p=0.017048) **(Figure 8c)**. Additionally, proteins found exclusively in PD-EV showed enrichment, in negative regulation of coagulation (p=0.000481), zymogen activation(p=0.000647), positive regulation of tau-protein kinase activity (p=0.00179), and acute-inflammatory response (p=0.00877) (**Figure 8e**). Molecular function showed enrichment in nitric-oxide synthase regulator activity (p=0.005538) and TPR domain binding (p=0.007748) (**Figure 8f**).

**Figure 8:**
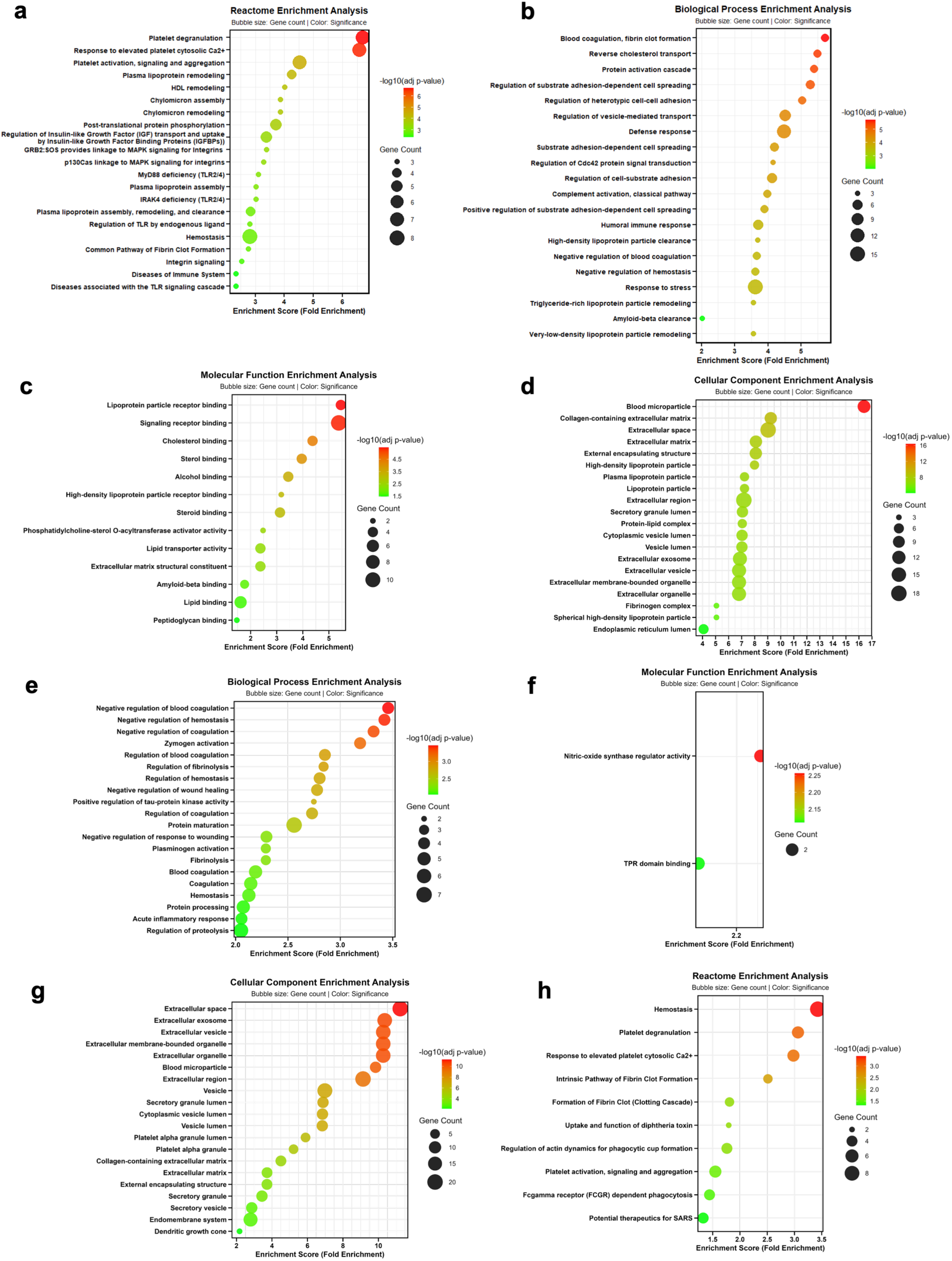
Gene ontology analysis. GO for downregulated proteins using enrichment bubble plots for **(a)** reactome pathway, **(b)** biological function, **(c)** molecular function, and **(d)** cellular component. **(e-h)** GO for proteins exclusively present in PD-EV. The x-axis represents enrichment score (-log_10_ of the adjusted p-value).

## Discussion

The present findings provide evidence that circulating EV derived from PD patients have direct neurotoxic effects on neuronal cells and induce a senescent, proinflammatory phenotype in microglia, which in turn exacerbates neuronal apoptosis **(Figure 9)**. We demonstrated that apoptosis induction was higher through the intermediation of microglia, whereas neurite toxicity was higher with direct PD-EV treatment. Previous studies have shown that PD-EV modulates microglial activity in murine models^57–60^ and elicits both morphological and functional changes in microglia by promoting a pro-inflammatory response^61,62^. Our findings extend these results and demonstrate that PD-EV triggers a complex activation pattern in HMC3 cells, characterized by increased levels of pro-inflammatory and senescence mediators. In fact, upon PD-EV treatment, we observed higher secretion of TNF-α, IFN-γ, IL-1β, and IL-8, which are classical pro-inflammatory SASP, and pro-apoptotic cytokines, as well as known markers of microglial activation^63–65^. Although it is challenging to distinguish between pro-inflammatory and senescent microglia due to the overlapping phenotypes, our results show that PD-EV caused up-regulation of IL-8 and IL-1β, which have been shown to be specifically elevated upon chemically-induction of senescence in HMC3 cells^66,67^. Classical senescence markers, including p21^WAF1/CIP1^ and p16^INK4a,^ have been reported in microglia within the aged mouse brain^68^. Despite the use of immortalized cells, which might present interference with the expression of cell-cycle related genes, we demonstrated that HMC3 cells showed transient induction of senescence-like phenotype after PD-EV treatment and displayed a trend towards increased expression of the senescence marker p16^INK4a^.

**Figure 9:**
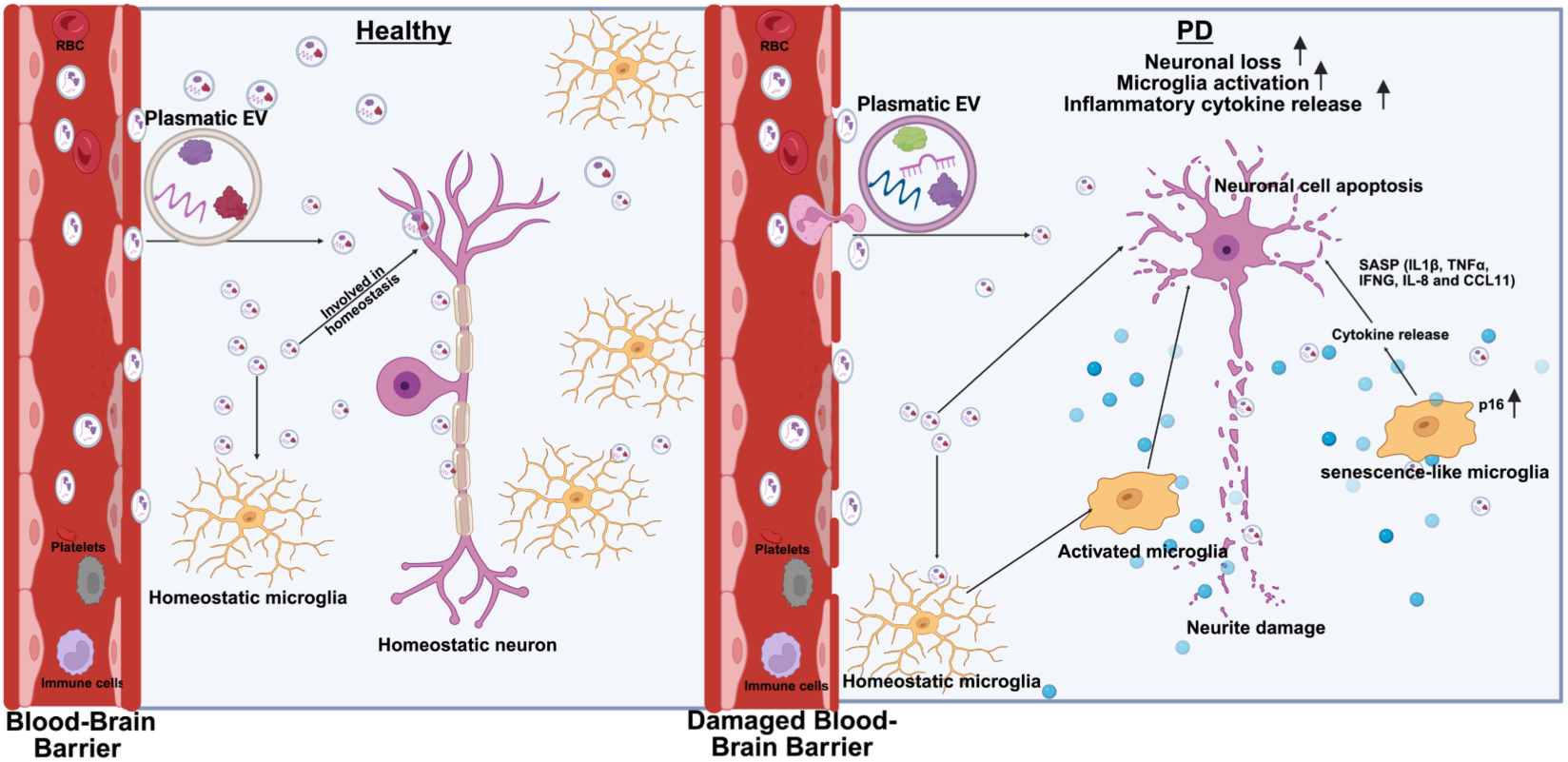
Involvement of plasmatic EV in PD neuroinflammation and neurodegeneration. Plasmatic EV from PD patients induce apoptosis in neurons. Senescence-like microglia promote neurotoxicity and exacerbate neuroinflammation via pro-inflammatory response, resulting in neuronal degeneration. Created with BioRender.

Since the largest risk factor for PD is ageing, senescent cells increase with age and chronic SASP is believed to be a key-driver for many diseases including neurodegenerative ones, our results highlight the relevance of circulating EV as possible targets for reversing or reducing brain cell senescence^69^.

We observed increased levels of CCL11, CCL17, TIMP-1 and GM-CSF in microglia after PD-EV treatment, which are involved in aging-related inflammation and alternative immune activation^70–72^. CCL11 plays an important role in modulating the adaptive immune response by promoting T cell-dependent neuroinflammation and nigrostriatal degeneration in patients with PD^73^. The elevated TIMP1 levels, along with reduced MMP1 and MMP2 levels, are consistent with previous studies on postmortem PD brains, which demonstrated decreased MMP2 and increased TIMP1 levels in the substantia nigra of patients with PD, suggesting that alterations in the MMP/TIMP system are linked to extracellular remodeling and disease pathogenesis^74^. Remarkably, we have shown increased levels of S100B in HMC3 treated with PD-EV: extracellular S100B at high concentration acts as a Damage-Associated Molecular Pattern (DAMP) protein, activates NF-κB-dependent transcription in microglia, which exhibit pro-inflammatory phenotypes^56^. It is conceivable that chronic DAMP exposure through paracrine or autocrine mechanisms locks microglia into persistent inflammatory activation and sustained stress signaling, leading to cell-cycle arrest and senescence with a pro-inflammatory SASP.

The elevated BDNF level in the secretome of HMC3 cells after PD-EV treatment is in line with previous studies showing that BDNF secretion from activated microglia likely promotes microglial proliferation into a proinflammatory state^75,76^. By functional pathway enrichment analysis of enriched cytokines, IL-17 signaling pathway emerged as the most prominent. IL-17 is an important effector of mucosal-associated invariant T (MAIT) and invariant natural killer T (iNKT) cells, which are enriched in mucosal organs and are reliable markers for compromised barrier integrity, for example, in gut^77^, which is in turn maybe a possible starting point for PD-related pathological cascade^78^.

Proteomic profiling of plasma EV from PD patients revealed the enrichment of complement activators, including C1QC, C5, CFH, and C4, in PD-EV, indicating active involvement of the complement cascade. This cascade activates microglial pro-inflammatory responses and abnormal synaptic pruning, consistent with our observation of reduced neurite length in PD-EV and increased secretion of TNF-α, IFN-γ, and IL-1β by microglia^79,80^. Increased levels of acute-phase proteins, such as SERPINs (SERPINA1), CD5L, and HP, as well as immunoglobulin fragments, which modulate immune response and cytokine production, indicate the inflammatory state of PD-EV^81,82^. Cofilin 1, a cytoskeleton regulator known to impair neuronal morphology and promote the seeding and spreading of alpha-synuclein aggregates^83^, is increased in PD-EV. Notably, the higher presence of cofilin 1 and calmodulin in PD-EV may suggest a contribution to the disorganization of the neuronal cytoskeleton, and this is supported by our observation of reduced neurite length and increased neuronal apoptosis in SH-SY5Y neurons after PD-EV treatment. Additional proteins, such as THBS1 and QSOX1, suggest altered ER stress regulation, which has been observed in patients with PD^84,85^. HSP90AA1, CLIC1, GSN, and SERPINF2 were exclusively found in PD-EV. HSP90AA1 and CLIC1 levels are higher in pro-inflammatory microglia^86,87^, whereas GSN is found in Lewy bodies and alpha-synuclein inclusions in PD^88^, confirming the pathological role of PD-EV. Of interest HSP90, functions as a DAMP and engages with pattern recognition receptors such as Toll-like receptor 4, CD91, and LOX - 1 to modulate immune and inflammatory responses and it has been reported in peripheral blood leukocytes of patients with Alzheimer’s disease^89^. Conversely, HC-EV contains PGLYRP2, a peptidoglycan recognition pattern involved in preventing a persistent inflammatory state^90^. The reduced expression in PD-EV of CLU, APOE, and APOA1, extracellular chaperon proteins, which participate in amyloid beta binding and clearance^91–93^, as well as of tau^94^ and alpha-synuclein fibrils^95,96^ is of high interest and worthing of further investigation. In fact, others have shown that CLU, a glycoprotein with several isoforms and involved in diverse complex functions, is reduced in plasmatic neuronal EV in PD^95,96^

This study has some limitations. First, the use of simple cell models, such as SH-SY5Y and HMC3 cells, which do not fully recapitulate the human brain, limits our ability to study the effects of EV on other relevant brain-resident cells, such as astrocytes. Future research should focus on more physiologically relevant human-specific cell models, such as 3D brain organoids, which better mimic the human brain, to elucidate the contribution of EV in PD initiation and progression. Second, the effects of circulating EV on alpha-synuclein aggregation, which might provide additional mechanistic insights, were not analyzed in the current study. Third, HMC3 cells are immortalized and replicative, and do not fully reflect the characteristics of primary senescent microglia. Therefore, future studies using iPSC-derived microglia or primary microglia are necessary to confirm whether plasmatic EV from PD patients promote sustained senescence-like features in PD.

In conclusion, these results offer new insights into the biological effects of plasmatic PD-EV, highlighting their crucial role in bridging peripheral immune cells with brain-resident cells. PD-EV induces senescent-like features and an inflammatory state in microglia which promotes neuronal toxicity. Proteomic analysis of plasma EV provided a detailed molecular framework in support of the role of PD-EV in immune processes, senescence and chronic inflammation. Understanding the underlying cellular mechanisms by which PD-EV modulates microglial activity and their role in neuronal dysfunction offers valuable insights into the development of novel therapeutic targets for PD.

## Abbreviations

EV: extracellular vesicles
PD: Parkinson’s Disease
HC: Healthy control
CM: Conditioned media
PD-EV-HMC3: PD-EV-treated HMC3
HC-EV-HMC3: HC-EV-treated HMC3
EV^-^ CM: EV-depleted conditioned media
GO: Gene Ontology
NTA: Nanoparticle tracking analysis
TEM: Transmission electron microscope
SEC: Size exclusion chromatography
RT: Room temperature
TFA: Trifluoroacetic acid
ACN: Acetonitrile
ABC: Ammonium bicarbonate
TOF: Time of flight
HPLC: High-performance liquid chromatography

## Acknowledgements

We would like to express our deep gratitude to all the subjects who participated to the NSIPD001 study and provided the biological material needed to conduct this research study.

## Author contributions

A.Y. performed the experiments, analyzed data, and drafted the manuscript. E.V. and S.P. contributed to EV isolation, and cell culture experiments and analysis. E.L. contributed to senescence experiments and analysis. M.P. performed LC-MS/MS analysis. A.R. performed TEM analysis. L.B., E.V. EL and A.K.L. reviewed and edited the manuscript. G.M. conceived and supervised the project, reviewed and drafted the manuscript. All authors reviewed and approved the final version of the manuscript.

## Ethical approval and consent to participate

This study was performed in line with the principles of the Declaration of Helsinki. Subjects were included according to the study protocol, approved by the Cantonal Ethics Committee (CETi 2895). All enrolled subjects gave written informed consent to the study.

## Data availability

The mass spectrometry proteomics data have been deposited to the ProteomeXchange Consortium via the PRIDE partner repository with the dataset identifier PXD075417. All quantified proteins are made available as supplementary datasets.

## Supplementary figures

**Figure S1:**
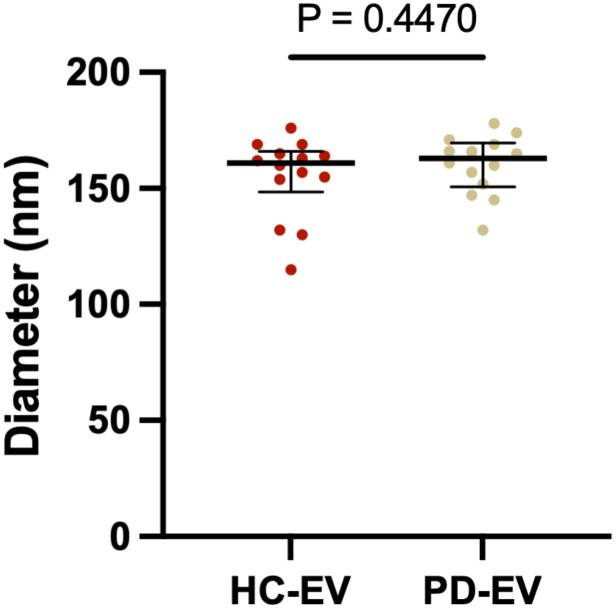
EV diameter from HC and PD. The graph presents all the data points, with the median represented as a straight line and IQR as an error bar. P-value is shown in the figure

**Figure S2:**
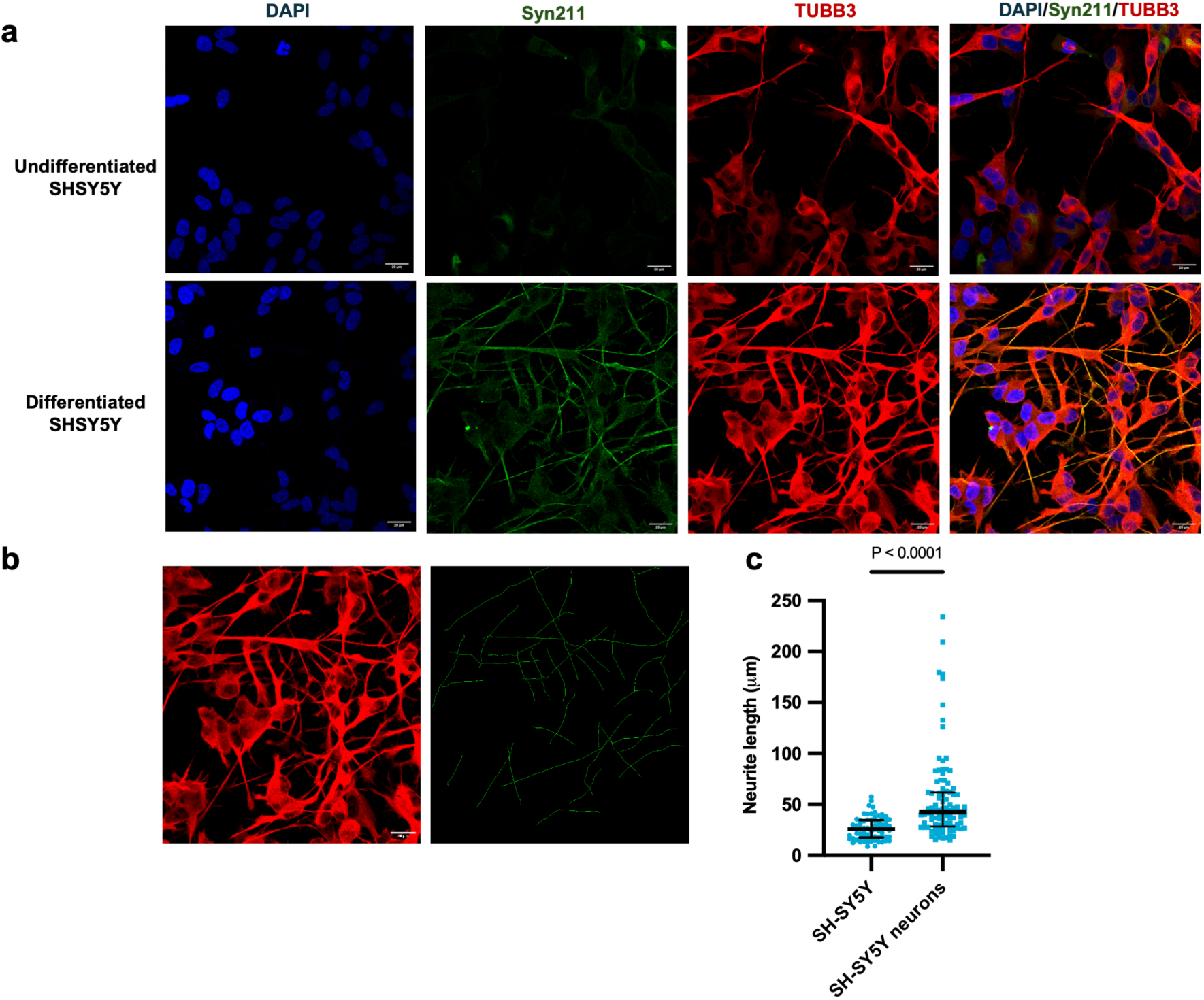
SHSY-5Y neurons differentiation. **(a-b)** Immunostaining of undifferentiated and differentiated SH-SY5Y with DAPI (blue), Syn211 (green) and TUBB3 (red). Scale bar: 20 µm. **(c)** Neurite length quantification of undifferentiated and differentiated SH-SY5Y (n=80-100 neurons per group). The graph shows all data points, with the median as a straight line and the IQR as an error bar for neurite length. P-value is shown in the figure

**Figure S3:**
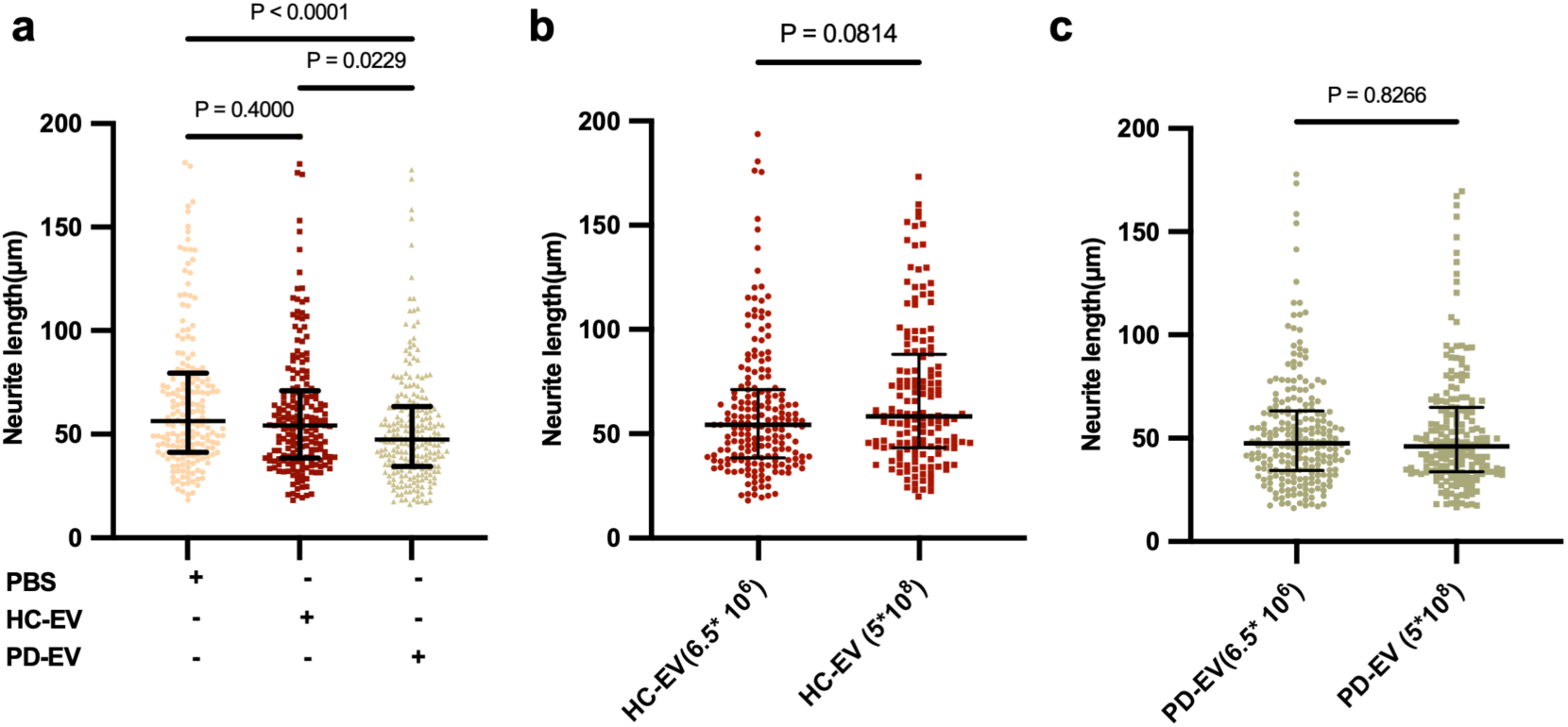
Neurite length reduction in SHSY5Y neurons. **(a)** SH-SY5Y neurite length quantification after treatment with 6.5 × 10^6^ EV/mL of PD-EV, HC-EV, and PBS (n=200-230 neurons per group). **(b-c)** Comparison of SH-SY5Y neurite length quantification after treatment with 6.5 × 10^6^ EV/mL or 5 × 10^8^ EV/mL of HC-EV and PD-EV. The graph presents all the data points, with the median represented as a straight line and the IQR as an error bar. P-values are shown in the figures

**Figure S4:**
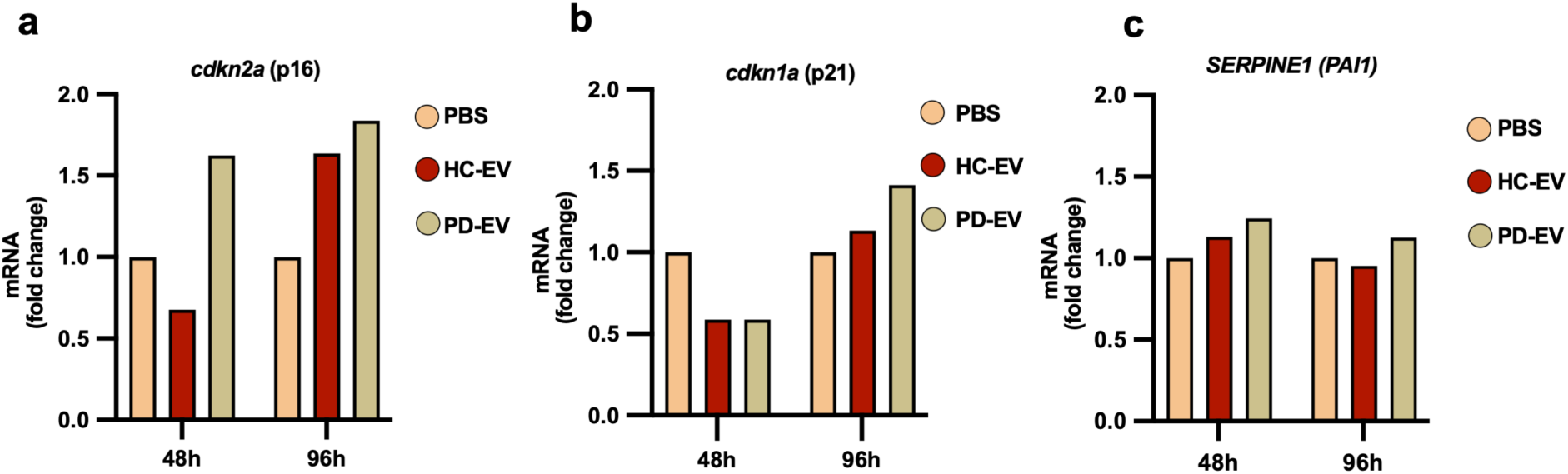
Expression of senescence-associated markers in HMC3 cells. **(a-c)** p16^INK4a,^ p21^WAF1/CIP1^, and SERPINE1 mRNA relative expression in HMC3 cells after 48 and 96 hours of treatment with HC-EV, PD-EV, or PBS.

**Figure S5:**
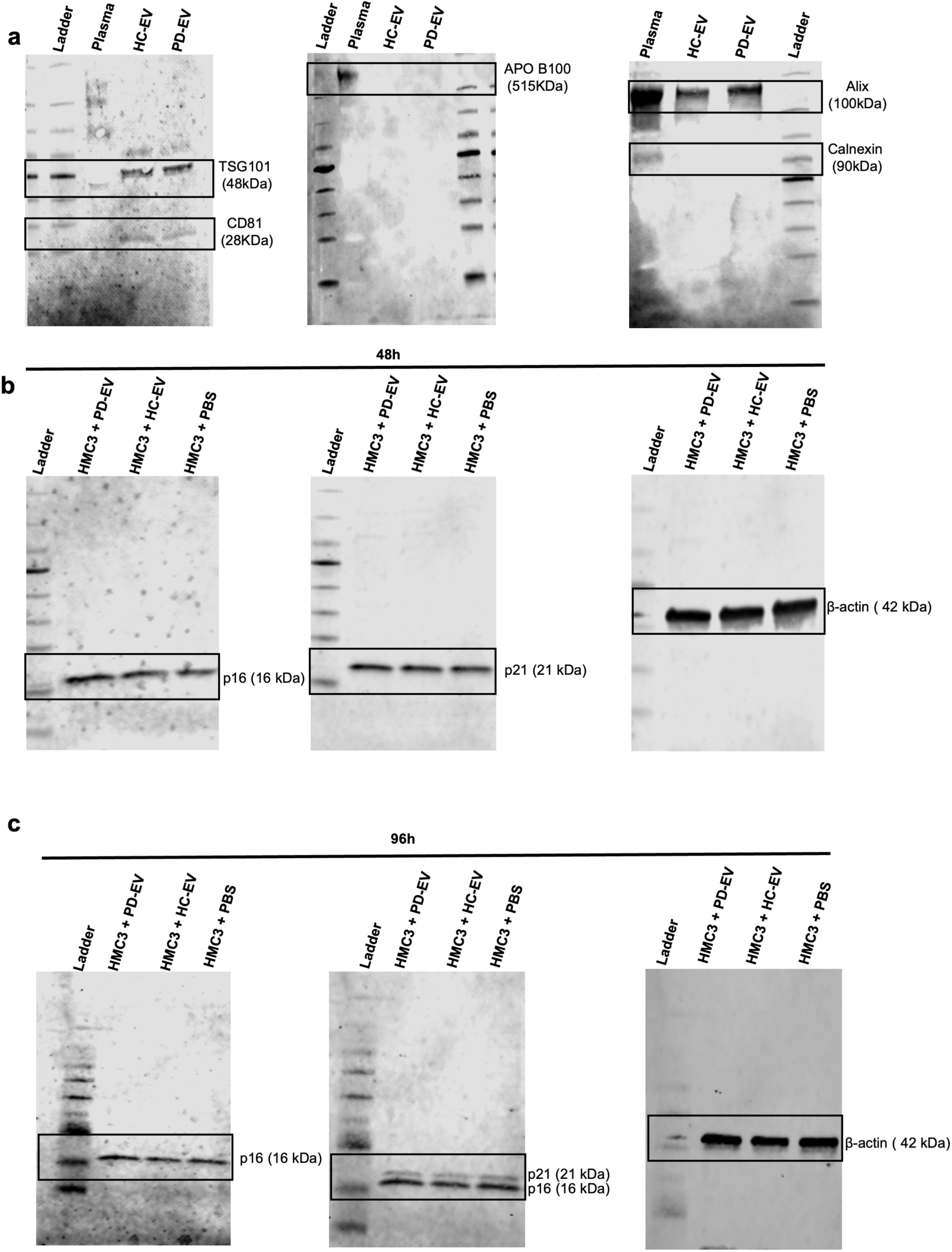
Uncropped western blot. **(a)** PVDF membrane blotted with anti-APO B100, anti-TSG101, anti-Alix, anti-Calnexin, and anti-CD81 antibodies. Black boxes represent the cropped images shown in Figure 1e. **(b-c)** PVDF membranes blotted with anti-p16, anti-p21, and anti-β-actin antibodies. The black box represents the cropped images shown in Figure 4a.

## References

1. Qin XY, Zhang SP, Cao C, Loh YP, Cheng Y. Aberrations in Peripheral Inflammatory Cytokine Levels in Parkinson Disease: A Systematic Review and Meta-analysis. JAMA Neurol. 2016;73(11):1316. doi:10.1001/jamaneurol.2016.2742

2. Poewe W, Seppi K, Tanner CM, et al. Parkinson disease. Nat Rev Dis Primer. 2017;3(1):17013. doi:10.1038/nrdp.2017.13

3. Jankovic J. Parkinson’s disease: clinical features and diagnosis. J Neurol Neurosurg Psychiatry. 2008;79(4):368–376. doi:10.1136/jnnp.2007.131045

4. Herman S, Djaldetti R, Mollenhauer B, Offen D. CSF-derived extracellular vesicles from patients with Parkinson’s disease induce symptoms and pathology. Brain. 2023;146(1):209–224. doi:10.1093/brain/awac261

5. Menšíková K, Matěj R, Colosimo C, et al. Lewy body disease or diseases with Lewy bodies? Npj Park Dis. 2022;8(1):3. doi:10.1038/s41531-021-00273-9

6. Kordower JH, Chu Y, Hauser RA, Freeman TB, Olanow CW. Lewy body–like pathology in long-term embryonic nigral transplants in Parkinson’s disease. Nat Med. 2008;14(5):504–506. doi:10.1038/nm1747

7. Li JY, Englund E, Holton JL, et al. Lewy bodies in grafted neurons in subjects with Parkinson’s disease suggest host-to-graft disease propagation. Nat Med. 2008;14(5):501–503. doi:10.1038/nm1746

8. Danzer KM, Krebs SK, Wolff M, Birk G, Hengerer B. Seeding induced by α-synuclein oligomers provides evidence for spreading of α-synuclein pathology. J Neurochem. 2009;111(1):192–203. doi:10.1111/j.1471-4159.2009.06324.x

9. Hansen C, Angot E, Bergström AL, et al. α-Synuclein propagates from mouse brain to grafted dopaminergic neurons and seeds aggregation in cultured human cells. J Clin Invest. 2011;121(2):715–725. doi:10.1172/JCI43366

10. Lauritsen J, Romero-Ramos M. The systemic immune response in Parkinson’s disease: focus on the peripheral immune component. Trends Neurosci. 2023;46(10):863–878. doi:10.1016/j.tins.2023.07.005

11. Horsager J, Andersen KB, Knudsen K, et al. Brain-first versus body-first Parkinson’s disease: a multimodal imaging case-control study. Brain. 2020;143(10):3077–3088. doi:10.1093/brain/awaa238

12. Terkelsen MH, Klaestrup IH, Hvingelby V, Lauritsen J, Pavese N, Romero-Ramos M. Neuroinflammation and Immune Changes in Prodromal Parkinson’s Disease and Other Synucleinopathies. Bloem BR, Brundin P, Tan EK, Harms A, Lindestam Arlehamn C, Williams-Gray C, eds. J Park Dis. 2022;12(s1):S149–S163. doi:10.3233/JPD-223245

13. Witoelar A, Jansen IE, Wang Y, et al. Genome-wide Pleiotropy Between Parkinson Disease and Autoimmune Diseases. JAMA Neurol. 2017;74(7):780. doi:10.1001/jamaneurol.2017.0469

14. Kang X, Ploner A, Wang Y, et al. Genetic overlap between Parkinson’s disease and inflammatory bowel disease. Brain Commun. 2022;5(1):fcad002. doi:10.1093/braincomms/fcad002

15. Drobny A, Ngo PA, Neurath MF, Zunke F, López-Posadas R. Molecular Communication Between Neuronal Networks and Intestinal Epithelial Cells in Gut Inflammation and Parkinson’s Disease. Front Med. 2021;8:655123. doi:10.3389/fmed.2021.655123

16. McGeer PL, Itagaki S, Boyes BE, McGeer EG. Reactive microglia are positive for HLA-DR in the substantia nigra of Parkinson’s and Alzheimer’s disease brains. Neurology. 1988;38(8):1285–1285. doi:10.1212/WNL.38.8.1285

17. Croisier E, Moran LB, Dexter DT, Pearce RK, Graeber MB. Microglial inflammation in the parkinsonian substantia nigra: relationship to alpha-synuclein deposition. J Neuroinflammation. 2005;2(1):14. doi:10.1186/1742-2094-2-14

18. Craig DW, Hutchins E, Violich I, et al. RNA sequencing of whole blood reveals early alterations in immune cells and gene expression in Parkinson’s disease. Nat Aging. 2021;1(8):734–747. doi:10.1038/s43587-021-00088-6

19. Harms AS, Thome AD, Yan Z, et al. Peripheral monocyte entry is required for alpha-Synuclein induced inflammation and Neurodegeneration in a model of Parkinson disease. Exp Neurol. 2018;300:179–187. doi:10.1016/j.expneurol.2017.11.010

20. Earls RH, Menees KB, Chung J, et al. NK cells clear α-synuclein and the depletion of NK cells exacerbates synuclein pathology in a mouse model of α-synucleinopathy. Proc Natl Acad Sci. 2020;117(3):1762–1771. doi:10.1073/pnas.1909110117

21. Kouli A, Camacho M, Allinson K, Williams-Gray CH. Neuroinflammation and protein pathology in Parkinson’s disease dementia. Acta Neuropathol Commun. 2020;8(1):211. doi:10.1186/s40478-020-01083-5

22. Mogi M, Harada M, Riederer P, Narabayashi H, Fujita K, Nagatsu T. Tumor necrosis factor-α (TNF-α) increases both in the brain and in the cerebrospinal fluid from parkinsonian patients. Neurosci Lett. 1994;165(1-2):208–210. doi:10.1016/0304-3940(94)90746-3

23. Mogi M, Harada M, Kondo T, et al. Interleukin-1β, interleukin-6, epidermal growth factor and transforming growth factor-α are elevated in the brain from parkinsonian patients. Neurosci Lett. 1994;180(2):147–150. doi:10.1016/0304-3940(94)90508-8

24. Chen X, Hu Y, Cao Z, Liu Q, Cheng Y. Cerebrospinal Fluid Inflammatory Cytokine Aberrations in Alzheimer’s Disease, Parkinson’s Disease and Amyotrophic Lateral Sclerosis: A Systematic Review and Meta-Analysis. Front Immunol. 2018;9:2122. doi:10.3389/fimmu.2018.02122

25. Welsh JA, Goberdhan DCI, O’Driscoll L, et al. Minimal information for studies of extracellular vesicles (MISEV2023): From basic to advanced approaches. J Extracell Vesicles. 2024;13(2):e12404. doi:10.1002/jev2.12404

26. Bellingham SA, Guo BB, Coleman BM, Hill AF. Exosomes: Vehicles for the Transfer of Toxic Proteins Associated with Neurodegenerative Diseases? Front Physiol. 2012;3. doi:10.3389/fphys.2012.00124

27. Denis HL, De Rus Jacquet A, Alpaugh M, et al. Erythrocyte-derived extracellular vesicles transcytose across the blood-brain barrier to induce Parkinson’s disease-like neurodegeneration. Fluids Barriers CNS. 2025;22(1):38. doi:10.1186/s12987-025-00646-9

28. Liu Z, Chan RB, Cai Z, et al. α-Synuclein-containing erythrocytic extracellular vesicles: essential contributors to hyperactivation of monocytes in Parkinson’s disease. J Neuroinflammation. 2022;19(1):53. doi:10.1186/s12974-022-02413-1

29. Currim F, Brown-Leung J, Syeda T, et al. Rotenone induced acute miRNA alterations in extracellular vesicles produce mitochondrial dysfunction and cell death. Npj Park Dis. 2025;11(1):59. doi:10.1038/s41531-025-00917-0

30. Li KL, Huang HY, Ren H, Yang XL. Role of exosomes in the pathogenesis of inflammation in Parkinson’s disease. Neural Regen Res. 2022;17(9):1898. doi:10.4103/1673-5374.335143

31. Counil H, Silva RO, Rabanel J, et al. Brain penetration of peripheral extracellular vesicles from Alzheimer’s patients and induction of microglia activation. J Extracell Biol. 2025;4(1):e70027. doi:10.1002/jex2.70027

32. Vacchi E, Burrello J, Di Silvestre D, et al. Immune profiling of plasma-derived extracellular vesicles identifies Parkinson disease. Neurol Neuroimmunol Neuroinflammation. 2020;7(6):e866.doi:10.1212/NXI.0000000000000866

33. Daniel SE, Lees AJ. Parkinson’s Disease Society Brain Bank, London: overview and research. J Neural Transm Suppl. 1993;39:165–172.

34. Vacchi E, Burrello J, Burrello A, et al. Profiling Inflammatory Extracellular Vesicles in Plasma and Cerebrospinal Fluid: An Optimized Diagnostic Model for Parkinson’s Disease. Biomedicines. 2021;9(3):230. doi:10.3390/biomedicines9030230

35. Forster JI, Köglsberger S, Trefois C, et al. Characterization of Differentiated SH-SY5Y as Neuronal Screening Model Reveals Increased Oxidative Vulnerability. SLAS Discov. 2016;21(5):496–509. doi:10.1177/1087057115625190

36. Taylor-Whiteley TR, Le Maitre CL, Duce JA, Dalton CF, Smith DP. Recapitulating Parkinson’s disease pathology in a three-dimensional human neural cell culture model. Dis Model Mech. 2019;12(4):dmm038042. doi:10.1242/dmm.038042

37. Korecka JA, Van Kesteren RE, Blaas E, et al. Phenotypic Characterization of Retinoic Acid Differentiated SH-SY5Y Cells by Transcriptional Profiling. Lim KL, ed. PLoS ONE. 2013;8(5):e63862. doi:10.1371/journal.pone.0063862

38. Kovalevich J, Langford D. Considerations for the Use of SH-SY5Y Neuroblastoma Cells in Neurobiology. In: Amini S, White MK, eds. Neuronal Cell Culture. Vol 1078. Methods in Molecular Biology. Humana Press; 2013:9–21. doi:10.1007/978-1-62703-640-5_2

39. Meijering E, Jacob M, Sarria J -C. F, Steiner P, Hirling H, Unser M. Design and validation of a tool for neurite tracing and analysis in fluorescence microscopy images. Cytometry A. 2004;58A(2):167–176. doi:10.1002/cyto.a.20022

40. Meijering E. Neuron tracing in perspective. Cytometry A. 2010;77A(7):693–704. doi:10.1002/cyto.a.20895

41. Wang YJ, Monteagudo A, Downey MA, Ashton-Rickardt PG, Elmaleh DR. Cromolyn inhibits the secretion of inflammatory cytokines by human microglia (HMC3). Sci Rep. 2021;11(1):8054. doi:10.1038/s41598-021-85702-8

42. Baek M, Yoo E, Choi HI, et al. The BET inhibitor attenuates the inflammatory response and cell migration in human microglial HMC3 cell line. Sci Rep. 2021;11(1):8828. doi:10.1038/s41598-021-87828-1

43. Livak KJ, Schmittgen TD. Analysis of Relative Gene Expression Data Using Real-Time Quantitative PCR and the 2−ΔΔCT Method. Methods. 2001;25(4):402–408. doi:10.1006/meth.2001.1262

44. Rappsilber J, Mann M, Ishihama Y. Protocol for micro-purification, enrichment, pre-fractionation and storage of peptides for proteomics using StageTips. Nat Protoc. 2007;2(8):1896–1906. doi:10.1038/nprot.2007.261

45. Cox J, Mann M. MaxQuant enables high peptide identification rates, individualized p.p.b.-range mass accuracies and proteome-wide protein quantification. Nat Biotechnol. 2008;26(12):1367–1372. doi:10.1038/nbt.1511

46. Cox J, Neuhauser N, Michalski A, Scheltema RA, Olsen JV, Mann M. Andromeda: A Peptide Search Engine Integrated into the MaxQuant Environment. J Proteome Res. 2011;10(4):1794–1805. doi:10.1021/pr101065j

47. Cox J, Hein MY, Luber CA, Paron I, Nagaraj N, Mann M. Accurate Proteome-wide Label-free Quantification by Delayed Normalization and Maximal Peptide Ratio Extraction, Termed MaxLFQ. Mol Cell Proteomics. 2014;13(9):2513–2526. doi:10.1074/mcp.M113.031591

48. Kim HJ, Rames MJ, Goncalves F, et al. Selective enrichment of plasma cell-free messenger RNA in cancer-associated extracellular vesicles. Commun Biol. 2023;6(1):885. doi:10.1038/s42003-023-05232-z

49. Reimand J, Isserlin R, Voisin V, et al. Pathway enrichment analysis and visualization of omics data using g:Profiler, GSEA, Cytoscape and EnrichmentMap. Nat Protoc. 2019;14(2):482–517. doi:10.1038/s41596-018-0103-9

50. Reimand J, Kull M, Peterson H, Hansen J, Vilo J. g:Profiler—a web-based toolset for functional profiling of gene lists from large-scale experiments. Nucleic Acids Res. 2007;35(suppl_2):W193–W200. doi:10.1093/nar/gkm226

51. Jamaly S, Ramberg C, Olsen R, et al. Impact of preanalytical conditions on plasma concentration and size distribution of extracellular vesicles using Nanoparticle Tracking Analysis. Sci Rep. 2018;8(1):17216. doi:10.1038/s41598-018-35401-8

52. Chun C, Smith AST, Kim H, et al. Astrocyte-derived extracellular vesicles enhance the survival and electrophysiological function of human cortical neurons in vitro. Biomaterials. 2021;271:120700. doi:10.1016/j.biomaterials.2021.120700

53. Prince JA, Oreland L. Staurosporine Differentiated Human SH-SY5Y Neuroblastoma Cultures Exhibit Transient Apoptosis and Trophic Factor Independence. Brain Res Bull. 1997;43(6):515–523. doi:10.1016/S0361-9230(97)00328-6

54. Zhang Z, Niu K, Huang T, et al. Microglia depletion reduces neurodegeneration and remodels extracellular matrix in a mouse Parkinson’s disease model triggered by α-synuclein overexpression. Npj Park Dis. 2025;11(1):15. doi:10.1038/s41531-024-00846-4

55. Basisty N, Kale A, Jeon OH, et al. A proteomic atlas of senescence-associated secretomes for aging biomarker development. Serrano M, ed. PLOS Biol. 2020;18(1):e3000599. doi:10.1371/journal.pbio.3000599

56. Michetti F, Clementi ME, Di Liddo R, et al. The S100B Protein: A Multifaceted Pathogenic Factor More Than a Biomarker. Int J Mol Sci. 2023;24(11):9605. doi:10.3390/ijms24119605

57. Xia Y, Zhang G, Kou L, et al. Reactive microglia enhance the transmission of exosomal α-synuclein via toll-like receptor 2. Brain. 2021;144(7):2024–2037. doi:10.1093/brain/awab122

58. Stuendl A, Kraus T, Chatterjee M, et al. Α-SYNUCLEIN in Plasma-Derived Extracellular Vesicles Is a Potential Biomarker of Parkinson’s Disease. Mov Disord. 2021;36(11):2508–2518. doi:10.1002/mds.28639

59. Herman S, Djaldetti R, Mollenhauer B, Offen D. CSF-derived extracellular vesicles from patients with Parkinson’s disease induce symptoms and pathology. Brain. 2023;146(1):209–224. doi:10.1093/brain/awac261

60. Guo M, Wang J, Zhao Y, et al. Microglial exosomes facilitate α-synuclein transmission in Parkinson’s disease. Brain. 2020;143(5):1476–1497. doi:10.1093/brain/awaa090

61. Chang C, Lang H, Geng N, Wang J, Li N, Wang X. Exosomes of BV-2 cells induced by alpha-synuclein: Important mediator of neurodegeneration in PD. Neurosci Lett. 2013;548:190–195. doi:10.1016/j.neulet.2013.06.009

62. La Torre ME, Panaro MA, Ruggiero M, et al. Extracellular Vesicles Cargo in Modulating Microglia Functional Responses. Biology. 2022;11(10):1426. doi:10.3390/biology11101426

63. Koprich JB, Reske-Nielsen C, Mithal P, Isacson O. Neuroinflammation mediated by IL-1β increases susceptibility of dopamine neurons to degeneration in an animal model of Parkinson’s disease. J Neuroinflammation. 2008;5(1):8. doi:10.1186/1742-2094-5-8

64. Brenner D, Blaser H, Mak TW. Regulation of tumour necrosis factor signalling: live or let die. Nat Rev Immunol. 2015;15(6):362–374. doi:10.1038/nri3834

65. Mount MP, Lira A, Grimes D, et al. Involvement of Interferon-γ in Microglial-Mediated Loss of Dopaminergic Neurons. J Neurosci. 2007;27(12):3328–3337. doi:10.1523/JNEUROSCI.5321-06.2007

66. Armanville S, Tocco C, Haj Mohamad Z, Clarke D, Robitaille R, Drouin-Ouellet J. Chemically Induced Senescence Prompts Functional Changes in Human Microglia-Like Cells. Wang R, ed. J Immunol Res. 2025;2025(1):3214633. doi:10.1155/jimr/3214633

67. García-Domínguez M. Interplay Between Aging and Glial Cell Dysfunction: Implications for CNS Health. Life. 2025;15(10):1498. doi:10.3390/life15101498

68. Hong B, Ohtake Y, Itokazu T, Yamashita T. Glial senescence enhances α-synuclein pathology owing to its insufficient clearance caused by autophagy dysfunction. Cell Death Discov. 2024;10(1):50. doi:10.1038/s41420-024-01816-8

69. Russo T, Riessland M. Age-Related Midbrain Inflammation and Senescence in Parkinson’s Disease. Front Aging Neurosci. 2022;14:917797. doi:10.3389/fnagi.2022.917797

70. Hoefer J, Luger M, Dal-Pont C, Culig Z, Schennach H, Jochberger S. The “Aging Factor” Eotaxin-1 (CCL11) Is Detectable in Transfusion Blood Products and Increases with the Donor’s Age. Front Aging Neurosci. 2017;9:402. doi:10.3389/fnagi.2017.00402

71. Zhang Y, Tang X, Wang Z, et al. The chemokine CCL17 is a novel therapeutic target for cardiovascular aging. Signal Transduct Target Ther. 2023;8(1):157. doi:10.1038/s41392-023-01363-1

72. Ishikawa J, Hirose H, Ishikawa S. Tissue Inhibitor of Matrix Metalloproteinase 1 Increases With Ageing and Can Be Associated With Stroke ― Nested Case-Control Study ―. Circ Rep. 2019;1(11):502–507. doi:10.1253/circrep.CR-19-0084

73. Chandra G, Roy A, Rangasamy SB, Pahan K. Induction of Adaptive Immunity Leads to Nigrostriatal Disease Progression in MPTP Mouse Model of Parkinson’s Disease. J Immunol. 2017;198(11):4312–4326. doi:10.4049/jimmunol.1700149

74. Lorenzl S, Albers DS, Narr S, Chirichigno J, Beal MF. Expression of MMP-2, MMP-9, and MMP-1 and Their Endogenous Counterregulators TIMP-1 and TIMP-2 in Postmortem Brain Tissue of Parkinson’s Disease. Exp Neurol. 2002;178(1):13–20. doi:10.1006/exnr.2002.8019

75. Gomes C, Ferreira R, George J, et al. Activation of microglial cells triggers a release of brain-derived neurotrophic factor (BDNF) inducing their proliferation in an adenosine A2A receptor-dependent manner: A2A receptor blockade prevents BDNF release and proliferation of microglia. J Neuroinflammation. 2013;10(1):780. doi:10.1186/1742-2094-10-16

76. Garcia-Contreras M, Thakor AS. Human adipose tissue-derived mesenchymal stem cells and their extracellular vesicles modulate lipopolysaccharide activated human microglia. Cell Death Discov. 2021;7(1):98. doi:10.1038/s41420-021-00471-7

77. Paiva RA, Salou M. MAIT and iNKT cells in tissue homeostasis and repair. Immunobiology. 2025;230(3):152917. doi:10.1016/j.imbio.2025.152917

78. Derkinderen P, Cossais F, Kulcsárová K, et al. How leaky is the gut in Parkinson’s disease? eBioMedicine. 2025;117:105796. doi:10.1016/j.ebiom.2025.105796

79. Gregersen E, Betzer C, Kim WS, et al. Alpha-synuclein activates the classical complement pathway and mediates complement-dependent cell toxicity. J Neuroinflammation. 2021;18(1):177. doi:10.1186/s12974-021-02225-9

80. Yao H, Tong W, Song Y, et al. Exercise training upregulates CD55 to suppress complement-mediated synaptic phagocytosis in Parkinson’s disease. J Neuroinflammation. 2024;21(1):246. doi:10.1186/s12974-024-03234-0

81. Halbgebauer S, Nagl M, Klafki H, et al. Modified serpinA1 as risk marker for Parkinson’s disease dementia: Analysis of baseline data. Sci Rep. 2016;6(1):26145. doi:10.1038/srep26145

82. Arredouani MS, Kasran A, Vanoirbeek JA, Berger FG, Baumann H, Ceuppens JL. Haptoglobin dampens endotoxin-induced inflammatory effects both *in vitro* and *in vivo*. Immunology. 2005;114(2):263–271. doi:10.1111/j.1365-2567.2004.02071.x

83. Yan M, Xiong M, Dai L, et al. Cofilin 1 promotes the pathogenicity and transmission of pathological α-synuclein in mouse models of Parkinson’s disease. Npj Park Dis. 2022;8(1):1. doi:10.1038/s41531-021-00272-w

84. Yao L, Lu F, Koc S, et al. LRRK2 Gly2019Ser Mutation Promotes ER Stress via Interacting with THBS1/TGF-β1 in Parkinson’s Disease. Adv Sci. 2023;10(30):2303711. doi:10.1002/advs.202303711

85. Morel C, Adami P, Musard JF, Duval D, Radom J, Jouvenot M. Involvement of sulfhydryl oxidase QSOX1 in the protection of cells against oxidative stress-induced apoptosis. Exp Cell Res. 2007;313(19):3971–3982. doi:10.1016/j.yexcr.2007.09.003

86. Smajić S, Prada-Medina CA, Landoulsi Z, et al. Single-cell sequencing of human midbrain reveals glial activation and a Parkinson-specific neuronal state. Brain. 2022;145(3):964–978. doi:10.1093/brain/awab446

87. Averaimo S, Milton RH, Duchen MR, Mazzanti M. Chloride intracellular channel 1 (CLIC1): Sensor and effector during oxidative stress. FEBS Lett. 2010;584(10):2076–2084. doi:10.1016/j.febslet.2010.02.073

88. Welander H, Bontha SV, Näsström T, et al. Gelsolin co-occurs with Lewy bodies in vivo and accelerates α-synuclein aggregation in vitro. Biochem Biophys Res Commun. 2011;412(1):32–38. doi:10.1016/j.bbrc.2011.07.027

89. Tukaj S. Hsp90 as a pathophysiological factor and emerging therapeutic target in atopic dermatitis. Front Immunol. 2025;16:1658399. doi:10.3389/fimmu.2025.1658399

90. Royet J, Gupta D, Dziarski R. Peptidoglycan recognition proteins: modulators of the microbiome and inflammation. Nat Rev Immunol. 2011;11(12):837–851. doi:10.1038/nri3089

91. Nelson AR, Sagare AP, Zlokovic BV. Role of clusterin in the brain vascular clearance of amyloid-β. Proc Natl Acad Sci. 2017;114(33):8681–8682. doi:10.1073/pnas.1711357114

92. DeMattos RB, Cirrito JR, Parsadanian M, et al. ApoE and Clusterin Cooperatively Suppress Aβ Levels and Deposition. Neuron. 2004;41(2):193–202. doi:10.1016/S0896-6273(03)00850-X

93. Cao S, Fu X, Li W, Wang P, Li C, Shang H. Protective role of apolipoprotein A and B in Parkinson’s disease: A prospective study from UK Biobank. Parkinsonism Relat Disord. 2025;132:107266. doi:10.1016/j.parkreldis.2025.107266

94. Wojtas AM, Carlomagno Y, Sens JP, et al. Clusterin ameliorates tau pathology in vivo by inhibiting fibril formation. Acta Neuropathol Commun. 2020;8(1):210. doi:10.1186/s40478-020-01079-1

95. Carini G, Mohammed S, Filippini A, Ramazzina I, Russo I. The Emerging Role of the Molecular Chaperone Clusterin in Parkinson’s Disease. Int J Mol Sci. 2025;26(13):6351. doi:10.3390/ijms26136351

96. Jiang C, Hopfner F, Katsikoudi A, et al. Serum neuronal exosomes predict and differentiate Parkinson’s disease from atypical parkinsonism. J Neurol Neurosurg Psychiatry. 2020;91(7):720–729. doi:10.1136/jnnp-2019-322588

